# A Computational Model of Hydrogen Peroxide Production in Liver and its Removal by Catalase and GSH-reliant Enzymes that Can Predict Intracellular H_2_O_2_ Concentration and Cell Death During Incidents of Extreme Oxidative Stress: (1) Applications to PBPK/PD Modeling of the Trivalent Arsenical DMA*^III^*, (2) Insights Obtained into (a) the Role of Critical GSH Depletion in Apoptosis and (b) How Intracellular H_2_O_2_ Concentration is So Tightly Regulated

**DOI:** 10.1101/2023.09.03.556145

**Authors:** L. M. Bilinsky

## Abstract

I present a simple computational model of H_2_O_2_ metabolism in hepatocytes and oxidative stress-induced hepatocyte death that is unique, among existing models of cellular H_2_O_2_ metabolism, in its ability to accurately model H_2_O_2_ dynamics during incidents of extreme oxidative stress such as occur in the toxicological setting. Versions of the model are presented for rat hepatocytes *in vitro* and mouse liver *in vivo*. This is the first model of cellular H_2_O_2_ metabolism to incorporate a detailed, realistic model of GSH synthesis from its component amino acids, achieved by incorporating a minimal version of Reed and coworkers’ pioneering model of GSH metabolism in liver. I demonstrate a generic procedure for coupling the model to an existing PK model for a xenobiotic causing oxidative stress in hepatocytes, using experimental data on hepatocyte mortality resulting from *in vitro* exposure to the xenobiotic at various concentrations. The result is a PBPK/PD model that predicts intracellular H_2_O_2_ concentration and oxidative stress-induced hepatocyte death; both *in vitro* and *in vivo* (liver of living animal) PBPK/PD models can be produced. I demonstrate the procedure for the ROS-generating trivalent arsenical DMA*^III^*. Simulations of DMA*^III^* exposure using the model indicate that critical GSH depletion is the immediate trigger for intracellular H_2_O_2_ rising to concentrations associated with apoptosis (*>* 1 *µM*), that this may only occur hours after intracellular DMA*^III^* peaks (“delay effect”), that when it does occur, H_2_O_2_ concentration rises rapidly in a sequence of two boundary layers, characterized by the kinetics of glutathione peroxidase (first boundary layer) and catalase (second boundary layer), and finally, that intracellular H_2_O_2_ concentration *>* 1 *µM* implies critical GSH depletion. Franco and coworkers have found that GSH depletion is central to apoptosis through mechanisms independent of ROS formation and have speculated that elevated ROS may simply indicate, rather than cause, an apoptotic milieu. Model simulations are consistent with this view, as they indicate that intracellular H_2_O_2_ concentration *>* 1 *µM* and extreme GSH depletion cooccur/imply each other; however, I note that this does not rule out a direct role for elevated ROS in the apoptotic mechanism. Finally, the delay effect is found to underlie a mechanism by which a normal-as-transient but pathological-as-baseline intracellular H_2_O_2_ concentration will eventually trigger critical GSH depletion and H_2_O_2_ concentration in the range associated with apoptosis, if and only if it persists for hours; this helps to rigorously explain how cells are able to maintain intracellular H_2_O_2_ concentration within such an extremely narrow range.

DISCLAIMER: The views presented in this article do not necessarily reflect those of the U.S. Food and Drug Administration or the National Toxicology Program.

## 1. Introduction

### 1.1. Reactive oxygen species (ROS); trivalent arsenicals

Reactive oxygen species (ROS) are byproducts of normal cellular metabolism, produced by mitochondria and by extramitochondrial cytoplasmic organelles including peroxisomes and endoplasmic reticulum (ER) [72]. The most abundant ROS in mammals are (1) superoxide anion (*·*O*^−^*), the primary ROS, formed by the one-electron reduction of molecular oxygen, (2) hydrogen peroxide (H_2_O_2_), produced from superoxide anion by the enzymes superoxide dismutase 2 (SOD2) in mitochondria and superoxide dismutase 1 (SOD1) in cytoplasm, and also produced in the peroxisomes by oxidizing enzymes, and (3) hydroxyl radical (*·*OH), produced when H_2_O_2_ reacts with Fe^2+^ in the Fenton reaction. ROS have been discovered to play an important role in cellular signaling, but are damaging to lipids, proteins, and nucleic acids when present at high enough concentrations, a situation referred to as one of “oxidative stress.”

Indeed, induction of oxidative stress is a generic mechanism by which many toxic xenobiotics cause harm. Among these are the trivalent arsenicals inorganic arsenite (iAs*^III^*), monomethylarsonous acid (MMA*^III^*), and dimethylarsinous acid (DMA*^III^*). Inorganic arsenic entering the liver is eliminated via an enzyme-catalyzed pathway consisting of a sequence of methylation steps which produce mono-, di-, and tri-methylated arsenicals [63]. In humans and rodents, the pentavalent form of the dimethylated metabolite (dimethylarsinic acid, DMA*^V^*) is the major elimination product appearing in urine and feces after ingestion or inhalation of arsenite or arsenate (pentavalent inorganic arsenic) [32]; inorganic arsenic and each of its methylated metabolites can exist in either a trivalent or a pentavalent chemical form and both are found inside cells, where they interconvert. It has been known for a long time that it is exclusively the trivalent forms which can undergo methylation, in a pathway catalyzed by arsenite methyltransferase (AS3MT). Although the trivalent arsenicals are necessary for elimination, it is now known that MMA*^III^* and DMA*^III^* are extremely cytotoxic and genotoxic [65, 43]. In *in vitro* experiments with rat hepatocytes, Naranmandura et al. determined that MMA*^III^* binds to targets in mitochondria and promotes ROS production by this organelle [43], while DMA*^III^* targets endoplasmic reticulum, elevating ROS production by that organelle [42]. Naranmandura et al. conducted experiments in which rat hepatocyte death was assayed after 24 hours’ exposure to DMA*^III^* in medium at concentrations ranging from 0.1 to 10 *µM* [43]. Significant cell death was found to occur for exposure levels at and above 1 *µM* DMA*^III^*. These results were compared with those for rat hepatocytes incubated for 24 hours in medium supplemented with 2000 *µM* N-acetyl cysteine (NAC, which is rapidly taken up by hepatocytes and converted to cysteine) prior to DMA*^III^* exposure. NAC was found to have a profoundly protective effect on cells, making the difference between approximately 80% survival (with NAC) and no survival (without NAC) after 24 hours’ exposure to 10 *µM* DMA*^III^*. NAC treatment supplies hepatocytes with a large amount of cysteine for use in the synthesis of reduced-form glutathione (GSH), a necessary cofactor for the GSH-reliant antioxidant defense system discussed shortly.

### 1.2. Need for a model of endogenous H_2_O_2_ production and removal in hepatocytes prior to constructing a model of xenobiotic-induced oxidative stress and resultant cell death; major innovations of the work presented here over existing models of cellular H_2_O_2_ metabolism

Given how generic a cytotoxic mechanism the generation of oxidative stress by a xenobiotic is, this phenomenon warrants a mathematical model. In this paper such a model is presented, for the case of hepatocytes exposed to DMA*^III^*, with aim of reproducing and rigorously explaining the experimental findings described above, due to Naranmandura and coworkers. Prior to constructing such a model, one must first model endogenous (that is, normal physiological) ROS production and removal in hepatocytes. This is because the consequences of DMA*^III^* for ROS load can only be modeled as a deviation/perturbation of this baseline situation.

As a first step, it is useful to create a model featuring just H_2_O_2_, since H_2_O_2_ is the major product of the primary ROS, superoxide anion, via the action of the superoxide dismutases. Under normal physiological circumstances, H_2_O_2_ is produced in liver at rate 50 nmol H_2_O_2_ per gram liver (wet weight) per minute, corresponding to 2-3% of total cellular oxygen uptake [59]. Under normal conditions, H_2_O_2_ is present at 0.01 *µM* in cytoplasm and 0.005 *µM* in mitochondria [8]. Intracellular concentrations *>* 1 *µM* are thought to trigger apoptosis [59] while extremely high concentrations cause necrosis [52]. H_2_O_2_ is reduced (hence removed) via three enzymatic systems: (1) Catalase (CAT), an enzyme which reduces two H_2_O_2_ molecules to two molecules of water and one molecule of molecular oxygen. In liver, CAT is mainly present in the peroxisomes of cytoplasm. In contrast to the other two H_2_O_2_-removal systems, CAT requires no cofactor that can become depleted to neutralize H_2_O_2_. However, the affinity of CAT for H_2_O_2_ is quite low (*K_m_ ∼* 10, 000 *µM* [69]) and only effects a quantitatively significant removal of H_2_O_2_ when H_2_O_2_ concentration is very high. (2) The GSH-based system, comprising the glutathione peroxidases and the glutaredoxins. Enzymes in this system, which reduce H_2_O_2_ itself or H_2_O_2_-oxidized proteins and lipids (containing organoperoxide and disulfide groups), become oxidized in the process and utilize the cofactor glutathione (GSH, gamma-L-glutamyl-L-cysteinyl-glycine), the most abundant non-protein source of thiol groups in cells, to return to their reduced, active enzymatic forms; as a result of this return to active form, GSH becomes oxidized. For example, glutathione peroxidase (GPX) catalyzes a reaction which consumes two GSH for each H_2_O_2_ removed, producing two molecules water and one molecule glutathione disulfide (GSSG), the oxidized, “spent” (unusable as a reductant) form of GSH. GSSG is recycled back to the active GSH form via the enzyme glutathione reductase (GR), which relies upon NADPH as electron donor. The cycling through the GSH and GSSG forms of glutathione by this antioxidant defense system is known as the glutathione redox cycle and is a major line of defense against oxidative stress. In liver, the organ in which most GSH synthesis occurs, GSH is normally present at about 10,000 *µM* [45, 51], and GSSG at about 100 *µM* [38, 48]. The ratio of GSH concentration to GSSG concentration is known as the glutathione redox ratio or glutathione redox balance and is often used as an index of oxidative stress in cells. It is about 100:1 in liver under normal circumstances; under conditions of oxidative stress, such as caused by dysfunction of the electron transport chain or exposure to certain toxins, the glutathione redox ratio falls to 10:1 or even 1:1 [12]. (3) The third means by which H_2_O_2_ (and H_2_O_2_-oxidized proteins and lipids) is reduced is the thioredoxin system, which acts in a manner analogous to the GSH-based system but uses the small protein thioredoxin as cofactor; thioredoxin is reduced back to its active form by thioredoxin reductase, which, like glutathione reductase, uses NADPH as final electron donor.

The endogenous production of H_2_O_2_ and its removal via these antioxidant systems is a matter of great physiological importance. Consequently, many computational models have been developed to describe H_2_O_2_ dynamics in isolated mitochondria and in whole cells of various types [16, 2, 31, 55, 60]; see also a recent work specific for liver cells [56]. Although these have yielded many important insights, the model presented here is uniquely equipped to accurately describe intracellular H_2_O_2_ dynamics in liver during episodes of extreme oxidative stress, such as occur in the toxicological setting. This is because previous models either do not feature GSH synthesis, treating GSH itself or the glutathione pool (GSH+2 GSSG) as a constant, or they assume that GSH is synthesized at a constant, unvarying rate. The most important innovation of the work I present here is that the kinetics of GSH synthesis are realistically modeled even during incidents of extreme oxidative stress, when GSH is predicted to be very low. This is accomplished by incorporating a simpified version of Reed and coworkers’ pioneering model for GSH metabolism in liver [50], which describes the formation of GSH from its three component amino acids in a sequence of two enzyme-catalyzed reactions: (1) glutamate-cysteine ligase (GCL), which produces the dipeptide gamma-L-glutamyl-L-cysteine, and (2) glutathione synthetase (GSS), which adds the glycine peptide. The first step, GCL, is rate-limiting and features the very important phenomenon of product inhibiton (one step removed) by GSH. This is a mechanism of metabolic control which causes GSH to be synthesized, when present at its normal physiological concentration of about 10,000 *µM*, at a rate balancing its rate of efflux from hepatocytes, but, when GSH is depleted, at a higher rate due to the lifting of the product inhibition on GCL.

There are two reasons this is an important innovation for modeling episodes of extreme oxidative stress: (1) The model presented here predicts that intracellular GSH can become extremely depleted when H_2_O_2_ concentration is elevated above its normal physiological value, such as occurs when the rate of H_2_O_2_ production exceeds its endogenous baseline, due to the presence of a xenobiotic such as DMA*^III^*, or due to intrinsic cellular pathology. It is important to model GSH dynamics accurately in the low-GSH regime, because the model predicts that, when it is very low, the concentration of GSH becomes crucially significant to hepatocytes’ ability to keep the intracellular H_2_O_2_ concentration under values associated with apoptosis, due to its impact on the kinetics governing how rapidly the GSH-reliant antioxidant system can clear H_2_O_2_. (2) From a toxicological standpoint, a model featuring GSH synthesis from its component amino acids, as well as the concentration dynamics of cysteine (see next paragraph), is necessary to simulate *in vitro* experiments in which the ability of NAC (a cysteine source for GSH synthesis) to reduce mortality in cells treated with ROS-generating xenobiotics is assessed, such as those performed by Naranmandura and coworkers for DMA*^III^* [43]. This data set was extremely useful for identifying the pharmacodynamic portion of the model, as it enabled me to infer (a) the relationship between intracellular DMA*^III^* concentration and the rate of production of H_2_O_2_ by extramitochondrial organelles, and (b) the dependence of cellular death rate on intracellular H_2_O_2_ concentration (as I discuss in *Conclusions*, there is some controversy regarding whether intracellular H_2_O_2_ concentration *>* 1 *µM* directly triggers apoptosis, or whether it simply indicates an apoptotic regime, the immediate cause being extreme GSH depletion; for simplicity, in the model presented here I assume that cellular death rate depends directly on intracellular H_2_O_2_ concentration). Analogous data sets (NAC/no-NAC cell mortality after a set number of hours, for various exposure levels) would be used in tandem with the *in vitro* computational model of endogenous H_2_O_2_ metabolism presented here, to construct PBPK/PD models for other xenobiotics known to generate oxidative stress in hepatocytes. I note that the parameter relating H_2_O_2_ concentration to cellular death rate should be roughly unchanged.

As mentioned earlier, in addition to catalase and the GSH-reliant system, the third leg of H_2_O_2_ removal is provided by the thioredoxin system. This system includes the peroxiredoxins (also known as the thioredoxin peroxidases), which are extremely efficient scavengers of H_2_O_2_ and thought to be responsible for the majority of peroxide clearance under normal conditions [17, 46, 30]. The goal of this work is to accurately model H_2_O_2_ dynamics under conditions of oxidative stress that are intense enough to cause cell death on a time scale of hours. Given this purpose, in this first model I neglect the action of the peroxiredoxins, viewing this as an acceptable first approximation. There are two reasons for this. (1) High H_2_O_2_ concentration is known to hyperoxidize some members of this enzyme family to a state that cannot be reduced by thioredoxin. It is speculated that this enables peroxiredoxins to serve as sensors of H_2_O_2_ which “shut off” when its concentration is elevated, triggering a local build-up of H_2_O_2_ that then oxidizes signaling proteins [46, 17, 44]. (2) Independent of the issue of hyperoxidation, I suspect that any clearance of H_2_O_2_ via the thioredoxin system will be quantitatively dominated by its clearance through GSH-based systems (glutathione peroxidases and glutaredoxins) during times of high oxidative stress, due to the ability of cells to rapidly synthesize *de novo* reduced-form, usable glutathione (GSH), but not reduced-form, usable thioredoxin. Such rapid *de novo* synthesis cannot occur for thioredoxin as it is a small protein containing about 100 amino acids and must be synthesized by ribosomes. This is reflected in Adimora and coworkers’ model of intracellular H_2_O_2_ dynamics, in which GSH and thioredoxin are synthesized at constant rates, with the rate of thioredoxin synthesis being set equal to a value approximately 1*/*1000 the rate of GSH synthesis [2]. For simplicity, I neglect it here.

### 1.3. Deliverables of this work

In the first deliverable of this work, I present an ordinary differential equation (ODE) model of endogenous H_2_O_2_ production and removal in rat hepatocytes in cell culture subsisting on Dulbecco’s Modified Eagle Medium (the cell culture medium used by Naranmandura et al. [43]), with no arsenical or other xenobiotic present. See *Methods* for a summary of the featured metabolic processes. Once constructed, I validate this *in vitro* computational model of endogenous H_2_O_2_ metabolism in hepatocytes by demonstrating that it makes the following predictions, consistent with empirical findings, without having been constructed for that purpose: (1) cellular stores of GSH can be depleted by even 95% without an immediate loss of viability [49], (2) under conditions of high oxidative stress, the glutathione redox ratio, GSH:GSSG, becomes *∼* 10 and even *∼* 1 [12].

I then modify my and coworkers’ 2019 *in vivo* PBPK model for oral or intravenous dosing of mouse with DMA*^III^* or DMA*^V^* [7] to create an *in vitro* version of this model. I conjoin it with the *in vitro* computational model of endogenous H_2_O_2_ production and removal in rat hepatocytes under the assumption, as proof of concept, that rat and mouse hepatocytes respond similarly to arsenicals, then use the combined model to simulate Naranmandura and coworkers’ 2011 experiments in which rat hepatocytes are exposed to DMA*^III^* [43]. Naranmandura et al. found that DMA*^III^* targets endoplasmic reticulum and increases the rate of ROS production by this organelle [42]. I model this phenomenon by multiplying the endogenous rate of production of H_2_O_2_ in cytoplasm (by extramitochondrial cytoplasmic organelles, including endoplasmic reticulum) by a monotonic-increasing function of DMA*^III^* concentration, of form 1 + *f* ([DMA*^III^*]). I identify *f* ([DMA*^III^*]), as well as a parameter *death* dictating cellular death rate as a function of intracellular H_2_O_2_ concentration, by fitting the model to their experimental data. This yields the second deliverable of this work, an *in vitro* PBPK/PD model for exposure of rat hepatocytes to DMA*^III^* in medium that predicts intracellular H_2_O_2_ concentration and oxidative stress-induced cell death. This model rigorously (i.e., quantitatively) explains why NAC had such a powerful protective effect on Naranmandura and coworkers’ cells.

Next, I modify the *in vitro* computational model of endogenous H_2_O_2_ production and removal in rat hepatocytes to construct an analogous *in vivo* model for mouse liver, incorporating known differences in enzyme kinetics between the species; this constitutes the third deliverable. Finally, I conjoin this to my and coworkers’ 2019 *in vivo* PBPK model for oral or intravenous dosing of mouse with DMA*^III^* or DMA*^V^* to produce an *in vivo* PBPK/PD model that predicts intracellular H_2_O_2_ concentration in mouse liver and hepatocyte death; this is the fourth deliverable. In [7] I predicted that orally administered DMA*^V^* is extensively reduced by mouse intestine to DMA*^III^*, the more toxic, trivalent form. This has since been proven experimentally [67]. Hence, a portion of an oral dose of DMA*^V^* enters the liver as DMA*^III^*. In [7] I augmented the PBPK model with a few simple equations to generate a proof-of-concept PBPK/PD model. I show that the PBPK/PD model developed here is far more realistic.

The *in vitro* and *in vivo* computational models presented here for endogenous H_2_O_2_ production and removal in hepatocytes are of generic value in toxicological modeling, as they can be conjoined with existing pharmacokinetic models for any xenobiotic known to cause oxidative stress in liver, to produce a PBPK/PD model predicting intracellular H_2_O_2_ concentration and oxidative stress-induced hepatocyte death, provided that *in vitro* cell mortality data sets analogous to those collected by Naranmandura et al. [43] for rat hepatocytes exposed to DMA*^III^* are available. The models also have value outside the toxicological setting, as they enable us to investigate the physiological consequences for hepatocytes of derangements in various components of endogenous H_2_O_2_ metabolism (e.g., abnormal values for the rates of production of H_2_O_2_ or GSH, or for the activities of antioxidant enzymes, etc.). Finally, and perhaps most importantly, this work provides theoretical support for the hypothesis [21, 22] that critical GSH depletion plays a direct role in the apoptotic mechanism (as opposed to solely an indirect role, by elevating H_2_O_2_ concentration), and uncovers a novel, GSH-reliant mechanism by which cells can differentiate between normal, transient elevations in H_2_O_2_ concentration, of use in signal transduction pathways, and persistent ones that are indicative of cellular pathology, the latter condition triggering apoptosis.

## 2. Methods

### 2.1. General information

Ordinary differential equations are used to model all processes. All dependent variables (aside from *N*, the fraction of living hepatocytes) are concentrations of biochemical sub-strates of interest in the modeled compartments. For example, [GSH*_cyt_*] is the concentration of GSH in cytoplasm, in units of micromoles per liter (*µM*). Each differential equation is an expression of mass-balance: the time rate of change in substrate concentration (units *µM/hr*) is the sum of the velocities of reactions producing the substrate or transporting it into the compartment, minus the sum of the velocities of reactions consuming the substrate or exporting it from the compartment, divided by compartment volume. Where possible, functional expressions for transport kinetics and for biochemical reaction kinetics, and associated parameter values, were taken from the published literature. Other parameter values were identified by requiring the model to have a steady-state solution (a special solution featuring concentrations which are unchanging in time; I note that all solutions, corresponding to different sets of initial concentrations at *t* = 0, rapidly converge to this special solution when no xenobiotic is present) at which concentrations of substrates in hepatocytes and in blood plasma (for the *in vivo* model) or medium (for the *in vitro* model), and reaction and transport fluxes (e.g., rate of efflux of GSH from liver or cultured hepatocytes to blood plasma or medium), are consistent with values reported in the literature. All fitting was done by hand, i.e., no optimizer was used to identify model parameters. Model equations were coded in MATLAB and solutions were computed using the ode15s solver (MATLAB R2020b, The MathWorks, Inc., Natick, MA, 2020). All model equations are provided and explained in the *Appendix*.

### 2.2. Modeling endogenous (normal physiological) H_2_O_2_ production and removal in hepatocytes: featured metabolic processes

The model is described in detail in the *Appendix* ; this section serves as a summary. Separate compartments are used for mitochondria and cytoplasm. I assume constant rates of production of H_2_O_2_ in the mitochondria compartment and in the cytoplasm compartment (by extramitochondrial cytoplasmic organelles, including endoplasmic reticulum), and a constant rate of efflux of H_2_O_2_ from the mitochondria compartment to the cytoplasm compartment. Values of these constants are inferred from the physiology literature; see *Appendix* for a detailed discussion of how these choices were made. I model the removal of H_2_O_2_ by catalase (cytoplasm compartment) and by the GSH-reliant antioxidant system (cytoplasm and mitochondria compartments). The model used here for GSH metabolism in hepato-cytes is a minimal version of the one published by Reed et al. [50]: one-carbon metabolism is not modeled, as it is not directly relevant, nor is the transulfuration pathway featured; instead, intracellular cysteine is assumed to be generated by the transulfuration pathway at constant rate 100 *µM/hr*. These two simplifications previously appeared in Bilinsky, Reed, and Nijhout’s 2015 work on acetaminophen overdose in rat [6], which incorporated a version of the GSH metabolism model. In the version of the GSH metabolism model utilized here, I have updated the kinetic expressions for the fluxes through GCL and GSS, based upon kinetic studies found in the physiology literature for rat (*in vitro* rat model) and mouse (*in vivo* mouse model). In addition, I have developed a new kinetic expression for the flux through cysteine dioxygenase to more accurately model the fate of intracellular cysteine when it is present at very high concentration (such as in the NAC experiments); see *Appendix* for details. Finally, I have added a mitochondria compartment to this version of the GSH metabolism model, modeling the exchange of GSH between mitochondria and cytoplasm (in accord with reports in the published literature), and modeling redox cycling between mitochondrial GSH and GSSG separately from redox cycling between cytoplasmic GSH and GSSG.

I briefly summarize the modeling of GSH metabolism in the current paper. Uptake by hepatocytes of cysteine in blood plasma (in the *in vivo* model) or in cellular medium (in the *in vitro* model) via the ASC transporter is modeled. The catabolism of cysteine via cysteine dioxygenase is modeled. The transporter-mediated export of GSH and GSSG from hepatocytes is modeled. For the *in vitro* model only, no breakdown of GSH outside of hepatocytes is modeled, as cellular medium lacks the kidney enzymes needed for this. The removal of H_2_O_2_ by GPX (consuming two molecules of the cofactor GSH for each H_2_O_2_ molecule reduced, and producing one molecule GSSG) and the reduction of GSSG back to GSH by GR (producing two molecules GSH for each molecule GSSG reduced) is modeled. I note that modeling H_2_O_2_ dynamics was not a goal of Reed and coworkers’ GSH metabolism model and H_2_O_2_ was treated there as a constant; this is why catalase was not modeled, which only becomes relevant at very high intracellular H_2_O_2_ concentrations. Here, I model catalase. I adjust the *V_max_* values for GPX and GR to reflect the removal of H_2_O_2_ under normal physiological conditions at rate matching its endogenous rate of production. In the work presented here, which serves as a stoichiometrically correct first model, the flux through glutathione peroxidase (GPX) represents all processes for removing H_2_O_2_ which ultimately rely upon the oxidation of GSH to be sustained (i.e., the GSH-reliant antioxidant system), whether this be directly (via GPX proper acting upon H_2_O_2_ itself) or indirectly (reduction of H_2_O_2_-oxidized proteins, by GPX proper in the case of organoperoxide groups and by glutaredoxin in the case of disulfide groups). Following the original model of Reed et al., intracellular glutamate and glycine concentrations are held constant at their normal physiological values.

Fig. 1 presents a schematic for the *in vitro* computational model of endogenous H_2_O_2_ production and removal in hepatocytes, and illustrates that intracellular DMA*^III^* (when present at nonzero concentration) elevates the rate of H_2_O_2_ production by endoplasmic reticulum. A schematic for the *in vivo* model is not included as it would be largely identical, with medium replaced by a liver capillary compartment and a blood plasma compartment. The *in vivo* model features an input of dietary cysteine to blood plasma and the breakdown of GSH into its component amino acids in blood plasma.

**Figure 1:**
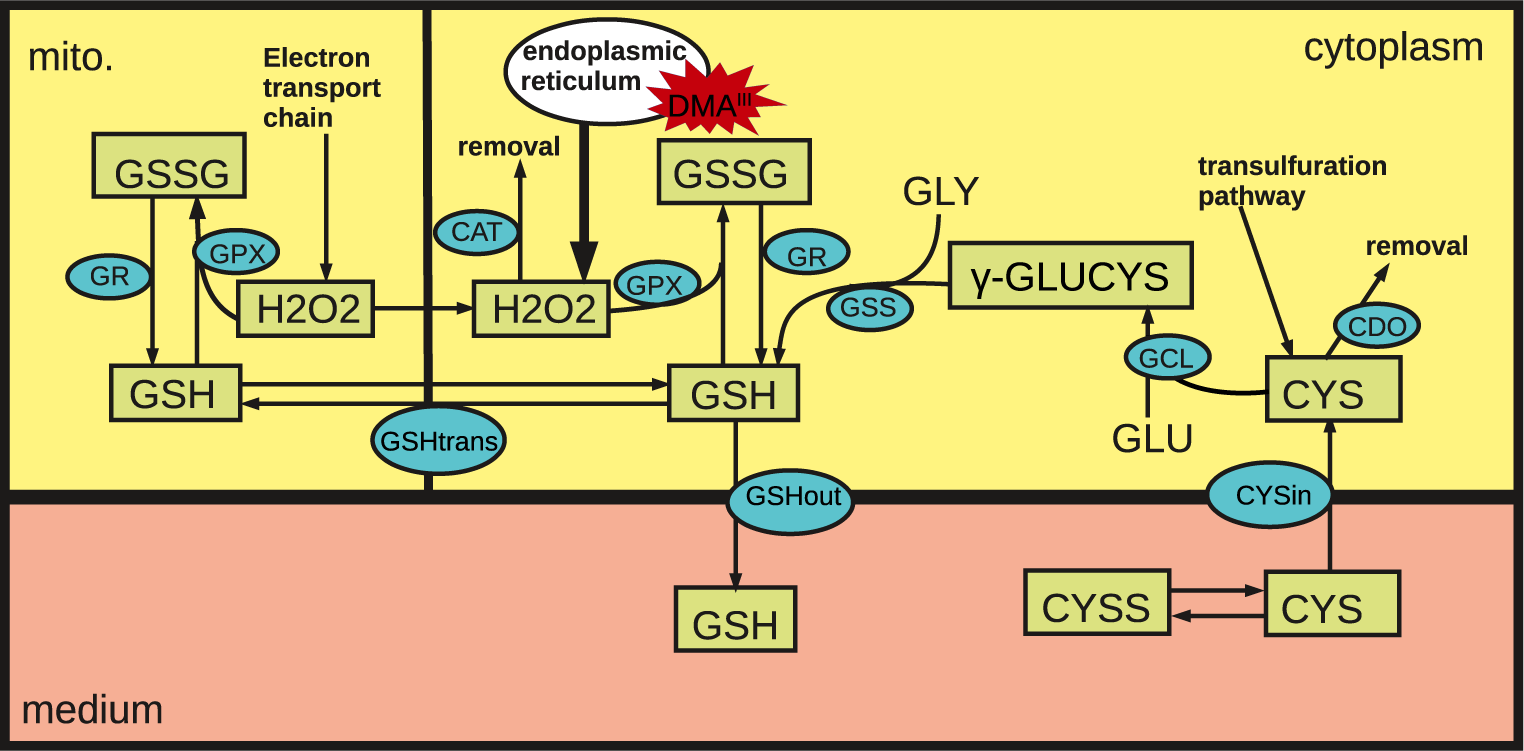
Schematic for the computational model of endogenous H_2_O_2_ production and removal in hepatocytes and the promotion of oxidative stress by DMA*^III^*. This particular schematic is for the *in vitro* rat hepatocyte model; the *in vivo* mouse liver model is similar. Modeled processes include the uptake by hepatocytes of cysteine from medium, the synthesis of GSH in cytoplasm, the loss of cysteine to catabolism by cysteine dioxygenase, the efflux of GSH from cells, the exchange of GSH between cytoplasm and mitochondria, the endogenous production of H_2_O_2_ in mitochondria by the electron transport chain, the diffusion of H_2_O_2_ from mitochondria to cytoplasm, the endogenous production of H_2_O_2_ in cytoplasm by organelles such as endoplasmic reticulum, the removal of H_2_O_2_ in cytoplasm by catalase, the removal of H_2_O_2_ in mitochondria and in cytoplasm by glutathione peroxidase, and the reduction of GSSG back to GSH in mitochondria and in cytoplasm by glutathione reductase. Intracellular DMA*^III^* increases the rate of production of ROS by endoplasmic reticulum [42], hence increasing the rate of H_2_O_2_ production in cytoplasm above baseline. Rectangles enclose the abbreviations of substrates whose concentrations are model variables; for each of these substrates, there is one differential equation modeling the time-evolution of its concentration. The concentrations of other substrates (glutamate and glycine) are held constant, at values equal to their normal physiological concentrations. Ellipses enclose the abbreviations of modeled enzymes and transporters. The production of H_2_O_2_ in mitochondria and in cytoplasm is not explicitly modeled; rather, constant rates for its production in mitochondria and in cytoplasm (when no DMA*^III^* is present), inferred from the physiology literature, are assumed. Similarly, the transulfuration pathway is not explicitly modeled but is assumed to result in a constant rate of appearance of intracellular cysteine in cytoplasm.

### 2.3. Modeling the induction of oxidative stress by xenobiotics and resultant hepatocyte death

The general strategy is to conjoin the *in vitro* computational model of endogenous H_2_O_2_ production and removal in hepatocytes with an *in vitro* PBPK model for an oxidative-stress-producing xenobiotic (here, DMA*^III^*, which interconverts in cells with its pentavalent counterpart DMA*^V^*), coupling the two models together via a function *f* which must be identified from experimental data reporting the degree of cell death caused by *in vitro* exposure of hepatocytes in culture to the xenobiotic; the data collected by Naranmandura et al. [43] reporting percentage cell death, by the 24-hour mark, for various exposure levels of DMA*^III^* in medium serves this purpose here. *f* appears in the differential equation for cytoplasmic H_2_O_2_ production (if I were modeling MMA*^III^*, which targets the mitochondria, it would appear in the equation for mitochondrial H_2_O_2_ production). It dictates the rate of H_2_O_2_ production in hepatocytes as a function of the intracellular xenobiotic concentration and is identified by using the conjoined model, which is an *in vitro* PBPK/PD model, to simulate the experiments that generated the cell mortality data; various definitions of *f* are tried until cell mortality at the 24-hour mark, as predicted by the model, agrees optimally with the experimental data. In this step the parameter *death* connecting oxidative stress to the cellular death rate is also identified; this parameter is not expected to be different for different xenobiotics (although it may differ between mouse and rat hepatocytes), as it is an intrinisic property of cells. Upon identification of *f*, one has an *in vitro* PBPK/PD model for the xenobiotic that predicts intracellular H_2_O_2_ concentration, and the fraction of living hepatocytes, as a function of time. I emphasize that different xenobiotics will generate different functions *f*.

Also provided in this work is an *in vivo* version of the computational model of endogenous H_2_O_2_ production and removal in hepatocytes, presented here for the case of mouse liver. To produce an *in vivo* PBPK/PD model for dosing of mouse with a xenobiotic that generates oxidative stress, this model would be conjoined with an extant *in vivo* PBPK model (which must include liver as one of the featured organs) for dosing of mouse with the xenobiotic, and the function *f*, identified in the course of developing, as a preliminary step, an *in vitro* PBPK/PD model (requiring mouse hepatocyte mortality data analogous to that collected by Naranmandura), inserted into the differential equation for either cytoplasmic or mitochondrial H_2_O_2_, depending on which organelle it targets. I demonstrate this procedure here for the case of DMA*^III^*, making use of my and colleagues’ 2019 *in vivo* PBPK model for oral or intravenous dosing of mouse with DMA*^III^* or DMA*^V^*. At the time of writing, no cell mortality data for mouse hepatocytes exposed to DMA*^III^* has been published analogous to that collected by Naranmandura et al. for rat hepatocytes. Hence, to generate the *in vivo* mouse PBPK/PD model, I assume that *f* is the same for mouse and rat hepatocytes. This is a rather large assumption and the mouse PBPK/PD model that results simply serves as a proof of concept.

Regarding the particular xenobiotic DMA*^III^* considered here, the pharmacokinetic modeling is rather complex. Trivalent and pentavalent arsenicals, including the dimethylated metabolites DMA*^III^* and DMA*^V^*, cannot (accurately) be separately quantified using analytic techniques. This makes it difficult to obtain the data sets needed to develop a PBPK model that is able to predict intracellular concentrations of DMA*^III^* and DMA*^V^* separately. To get around this obstacle, our 2019 PBPK model was constructed to predict a lower bound on the DMA*^III^* (the more acutely toxic valence form, which is the species responsible for the generation of oxidative stress) concentration and an upper bound on the DMA*^V^* concentration in liver and other organs, rather than these concentrations themselves, as the former could be done with greater accuracy. That is, model simulations enabled us to conclude “there is at least this much DMA*^III^* present, and at most this much DMA*^V^* present, at time *t*” in organs of interest after a given oral or intravenous dose of DMA*^III^* or DMA*^V^*. Analogous statements hold for the *in vitro* PBPK model derived in the *Appendix*, used to simulate Naranmandura and coworkers’ experiments. For ease of exposition, in this paper the computed lower bound on intracellular DMA*^III^* concentration, and upper bound on intracellular DMA*^V^* concentration, will be referred to as the computed intracellular DMA*^III^* and DMA*^V^* concentrations, respectively. This is equivalent to assuming the most extreme metabolic scenario, vis a vis intracellular removal of DMA*^III^* (which occurs via its oxidation to DMA*^V^* and subsequent efflux from cells), that is consistent with the PK data used to calibrate the model; see Bilinsky et al. [7].

## 3. Results and Discussion

### 3.1. in vitro computational model of endogenous H_2_O_2_ production and removal in rat hepatocytes in cell culture (no xenobiotic present)

Biochemical substrates in rat hepatocytes are present, under the model, at the concentrations given in Table 1 when no xenobiotic is present and the cells are subsisting on Dulbecco’s Modified Eagle Medium (DMEM). These concentrations are in agreement with the normal physiological values reported in the literature and the concentrations of amino acids in DMEM. This is the special steady-state solution discussed in the *Methods* section.

**Table 1:** Steady-state concentrations of biochemical substrates computed for the *in vitro* computational model of endogenous H_2_O_2_ production and removal in rat hepatocytes subsisting on Dulbecco’s Modified Eagle Medium (no xenobiotic present).

| Model concentration ($\mu M$) | Normal physiological range/comment | Source |
| --- | --- | --- |
| **Intracellular** |  |  |
| $H_2O_{2,cyt}=0.01$ | 0.01-0.1 | [8] |
| $H_2O_{2,mito}=0.005$ | 0.005 | [8] |
| $GSH_{cyt}=9438$ | 7631-12657 | [64] |
|  | 9643-10757 | [29] |
| $GSH_{mito}=9415$ | similar to cytoplasmic value | [51] |
| $GSSG_{cyt}=95$ ($\approx 2\%$ total GSH equivalents) | 0.24-1.9% (NEM method, whole liver) | [38] |
|  | 1-10.8% (2-VP method, whole liver) | [38] |
| $GSSG_{mito}=95$ ($\approx 2\%$ total GSH equivalents) | 1.25-2.44% (mitochondria of mouse) | [48] |
| γ-GC=9.93 | | |
| Cys=103 | 28-143 | [62] |
| **Medium** |  |  |
| Cys=20 | 8-40 (normal plasma concentration in rat) | [62],[53] |
| Cyss=250.5 | concentration of cystine in Dulbecco’s Modified Eagle Medium, less the cystine equivalents in 20 $\mu M$ cysteine (10 $\mu M$) | |

### 3.2. in vitro PBPK/PD model predicting H_2_O_2_ concentration in rat hepatocytes and cell death after exposure to the ROS-producing arsenical DMA^III^

In Naranmandura and coworkers’ experiments [43], rat hepatocytes in cell culture, subsisting on Dulbecco’s Modified Eagle Medium (DMEM), were exposed to various concentrations (0.1-10 *µM*) of DMA*^III^* for 24 hours, after which cell death was assayed and the percentage of living hepatocytes (relative to the starting number) computed. Prior to DMA*^III^* exposure, cells either had or had not (control group) been cultured, over the 24-hour period immediately preceding the addition of DMA*^III^* to medium, in DMEM supplemented with 2000 *µM* NAC. Their experimental data (solid circular markers) are reproduced in

Fig. 2. These data dramatically demonstrate the power of NAC to rescue cells from DMA*^III^*-induced toxicity, increasing the percentage of hepatocytes still alive after 24 hours’ exposure to DMA*^III^*. Simulation of these experiments enabled me to identify the function *f* defining the relationship between intracellular DMA*^III^* concentration and the rate at which H_2_O_2_ is produced in cytoplasm (that is, produced by extramitochondrial organelles such as endoplasmic reticulum), discussed in the next section.

**Figure 2:**
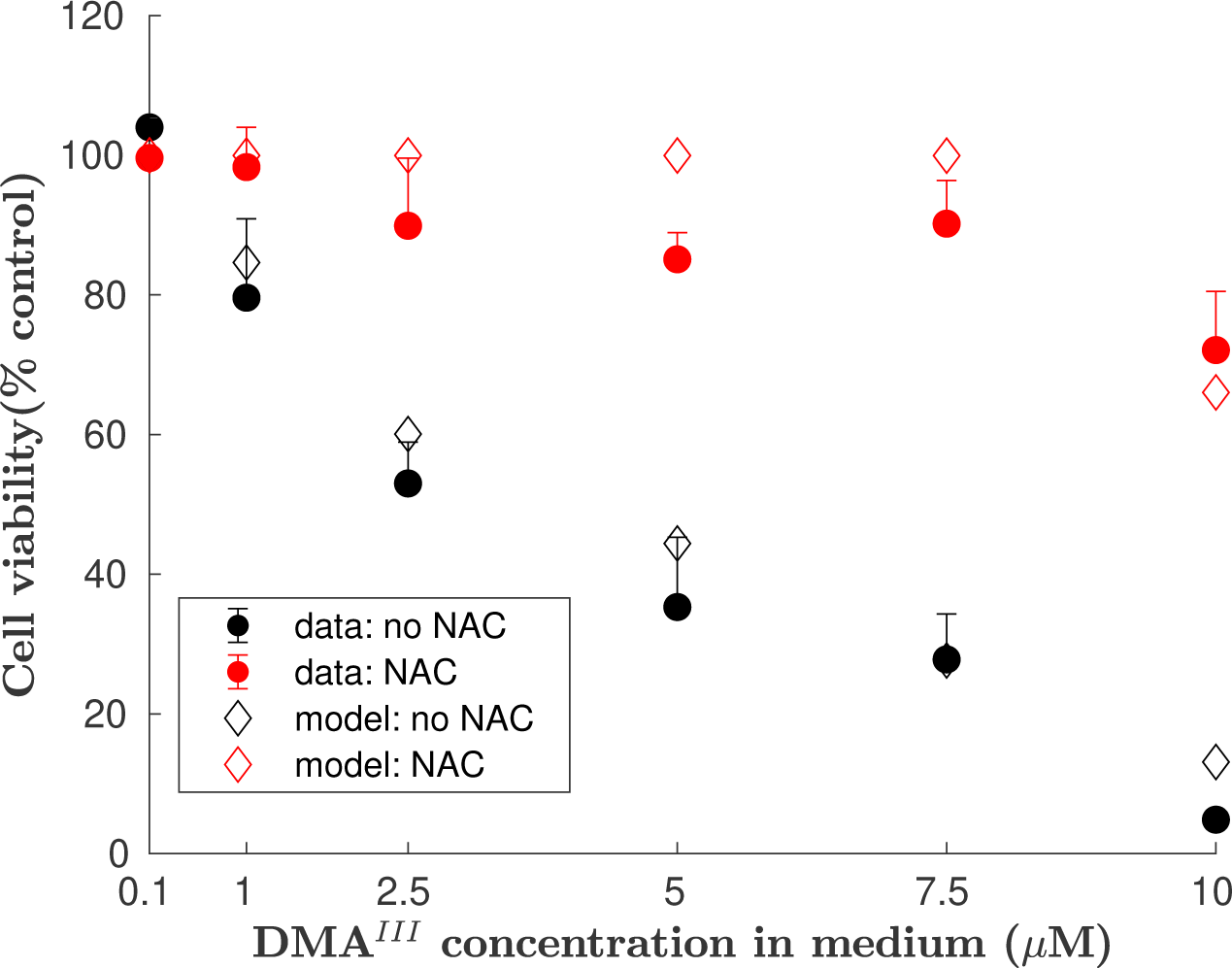
Simulation of Naranmandura experiments: DMA*^III^* -induced cell death by the 24-hour mark, with and without NAC supplementation. I simulate the experiments conducted by Naranmandura et al. [43] in which rat hepatocytes were exposed to various concentrations of DMA*^III^* in medium for 24 hours and cell viability assayed, with or without having been cultured for 24 hours in medium supplemented with 2000 *µM* NAC prior to DMA*^III^* exposure. The function *f* dictating the dependence of extramitochondrial H_2_O_2_ production upon intracellular DMA*^III^* concentration was identified by varying its functional form and parameter values until optimal agreement (shown in this figure) was achieved between model solutions and Naranmandura’s data. The fit is excellent: Average Absolute Fold Error= 1.17 and 10*^RMSLE^*=1.36 where RMSLE is the Root Mean Squared Logarithmic Error. Exposure of cells to 7.5 *µM* DMA*^III^* in medium without NAC supplementation results in approximately 70% cell death after 24 hours; with NAC supplementation, almost perfect rescue is observed. This is consistent with the NAC/no-NAC intracellular H_2_O_2_ concentrations predicted by the model, shown in Fig. 4.

In running a simulation, one must first define an initial condition from which the solution advances. For the control cell culture, this was defined as follows. The initial (*t* = 0, defined as the time at which DMA*^III^* is added to medium) concentrations of non-arsenical biochemical reactants in hepatocytes and in medium were set to the values given in Table 1, the steady state discussed in the previous section. The initial fraction of living hepatocytes was set to 1. The initial concentration of DMA*^III^* in medium was set to one of the values tested by Naranmandura et al. (0.1-10 *µM*), while the initial concentrations of intracellular DMA*^III^*, as well as those of both intracellular DMA*^V^* and DMA*^V^* in medium, were set to zero. This set of starting-point values for model dependent variables comprised the “initial condition” needed to run a simulation. Using this initial condition, the solution from *t* = 0 to *t* = 24 was computed, for a particular choice of the function *f*.

For the NAC-treated cell culture, the initial condition used to run a simulation was different, and computed as follows. A 24-hour “pre-simulation” (from *t* = *−*24 to *t* = 0) was run with no arsenicals present. The initial condition for this pre-simulation had biochemical reactant concentrations set to the values given in Table 1, except that the concentration of cysteine in medium was set to 2020 *µM* rather than 20 *µM*, reflecting the addition of 2000 *µM* NAC to medium. From this starting point the solution was advanced by 24 hours (from *t* = *−*24 to *t* = 0), simulating 24 hours’ pre-culturing with 2000 *µM* NAC. The end point of this pre-simulation provided the values of biochemical reactant concentrations appearing in the initial condition used for the actual simulation of the NAC-treated cells’ exposure to DMA*^III^* (from *t* = 0 to *t* = 24 hours); by comparison, these values were taken directly from Table 1 in defining the initial condition for the control cell culture. Aside from this, the initial condition used for the NAC-treated cell culture was the same as for the control cell culture: the fraction of living hepatocytes was set to 1, the concentration of DMA*^III^* in medium was set to a particular concentration tested by Naranmandura and coworkers, and the concentrations of intracellular DMA*^III^*, intracellular DMA*^V^*, and DMA*^V^* in medium were all set to 0. Using this initial condition, the solution from *t* = 0 to *t* = 24 was computed. The function *f* was assumed to be the same as for the control cell culture.

To summarize, for a given definition of the function *f*, the fraction of hepatocytes still alive by the 24-hr mark (*N* (24)), for each of the DMA*^III^* concentrations tested by Naranmandura and for both the control and NAC-treated cell cultures, was computed and plotted (hollow diamond markers) alongside Naranmandura’s experimental data. *f* was varied until optimal agreement, shown in Fig. 2, was obtained between the experimental data and model predictions for 24-hour cell viability.

I emphasize that the simulations of the control and NAC-treated cell cultures used to generate Fig. 2 differ only in the initial condition used at *t* = 0, when DMA*^III^* is introduced to medium. The initial condition used for the NAC-treated cell culture, computed as described above, featured a supraphysiological GSH concentration, due to the 24 hours of preculturing with NAC. It also featured a higher-than-normal concentration of cysteine in medium, as not all of the added 2000 *µM* NAC had been utilized by cells for GSH synthesis by the time DMA*^III^* exposure began.

#### 3.2.1. Simulation of Naranmandura experiments enables inference of the function f([DMA^III^]) relating intracellular DMA^III^ concentration to the rate of production of H_2_O_2_ in cytoplasm by extramitochondrial organelles

The first result to present is the formula for the function *f* ([DMA*^III^*]) relating the computed intracellular DMA*^III^* concentration to the rate of cytoplasmic H_2_O_2_ production (meaning, in the model, production by extramitochondrial organelles including endoplasmic reticulum; the production of H_2_O_2_ by mitochondria is modeled separately). *f* appears in the equation for cytoplasmic H_2_O_2_ concentration given below and causes the rate of cytoplasmic H_2_O_2_ production to exceed its normal endogenous value *h*2*o*2*prodcyt* when DMA*^III^* is present. See *Appendix* for a detailed explanation of this and all model equations.

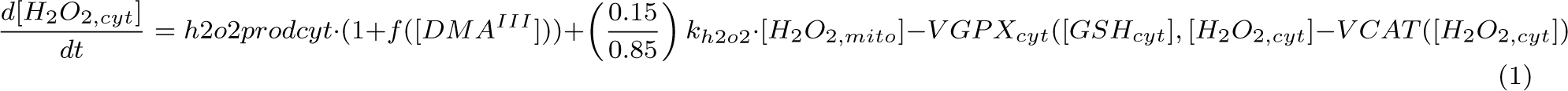

The identified optimal function *f* used to generate Fig. 2 is given in Eq. 2. In the process of identifying *f*, the parameter *death*, dictating the per-cell death rate of hepatocytes as a function of intracellular H_2_O_2_ concentration, was also identified and is given in Table A.4.

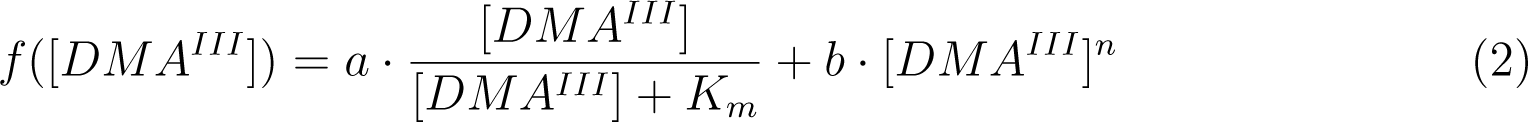

In this equation, *a* = 2.2, *K_m_* = 0.2 *µM*, *b* = 0.0005 *µM ^−^*^2.5^, and *n* = 2.5. DMA*^III^* is speculated to increase the rate at which ROS are produced by endoplasmic reticulum by binding to the thiol groups of proteins awaiting folding inside its lumen, blocking the formation of all necessary disulfide bridges of these proteins. Hence, these proteins do not undergo successful folding and export, but remain in the lumen, where futile redox cycling on their (non-DMA*^III^*-bound) thiol groups drives excessive ROS production [42]. I remark that any intracellular DMA*^III^* not so occupied would be expected to be bound elsewhere: DMA*^III^* and other trivalent arsenicals bind extensively to thiol groups of cellular proteins and also to GSH [19], and trivalent arsenicals are not expected to be present in free form to any significant extent in cells. In fact, the actual enzyme substrates for the methylation pathway by which arsenic is eliminated are the protein-bound or glutathione conjugates of the trivalent arsenicals [41, 25, 18]. I note that the function *f* identified here could be viewed as the composition of two functions which are not separately identifiable from this data set:

*f* = *g*(*h*([*DMA^III^*])), where *h* reflects the amount of DMA*^III^*-bound protein trapped inside the lumen, and *g* reflects the increase in ROS production due to futile redox cycling.

#### 3.2.2. Critical GSH depletion in non-NAC-treated cells triggers a rapid rise in intracellular H_2_O_2_ concentration to values in the range associated with apoptosis. H_2_O_2_ dynamics feature two boundary layer transitions associated, respectively, with (1) glutathione peroxidase kinetics and (2) catalase kinetics

When non-NAC-treated (control) hepatocytes were exposed to 7.5 *µM* DMA*^III^* in medium (starting concentration, at *t* = 0 *hr*), about 70% of cells died by the 24-hour mark. Simulations predict that these cells experienced a *>* 90% decline in GSH concentration, which was then sustained for the rest of the exposure period. This is despite the fact that under the model, (1) hepatocytes are at all times intensively synthesizing GSH from medium cysteine and (2) glutathione reductase (GR) is “recycling” spent glutathione (GSSG, the oxidized form) to the active, reduced form of GSH. The depletion in GSH is driven by the flux through glutathione peroxidase (GPX), representing in the model the GSH-reliant leg of cellular antioxidant defense. The presence of DMA*^III^* in hepatocytes elevates the rate of production of H_2_O_2_ in cytoplasm above its baseline value, according to the function *f* ([DMA*^III^*]). This effects an immediate, very mild increase in H_2_O_2_ concentration, to within an order of magnitude of its normal physiological concentration of 0.01 *µM* ; see Fig. 3. The slight elevation in H_2_O_2_ concentration drives the oxidation of GSH by GPX at rate slightly greater than what is normal, causing a gradual decrease in GSH concentration during what I call an “induction period.” During the induction period GSH is in decline and H_2_O_2_ concentration rising but not yet in the range associated with apoptosis. Until GSH has declined to concentrations for which the rate of removal of H_2_O_2_ via GPX (which is kinetically dependent upon both the GSH and H_2_O_2_ concentrations) is significantly less than the rate at which H_2_O_2_ is being produced, H_2_O_2_ concentration remains within tight bounds. It takes about an hour and a half for the mismatch between the rate of H_2_O_2_ production and the rate of removal to become significant, during which time H_2_O_2_ rises to about 0.1 *µM*, a level that has physiological uses but is pathological as a baseline [59]. See insets of Fig. 3. At about *t* = 1.44 *hr*, the mismatch becomes significant and H_2_O_2_ concentration climbs by an order of magnitude within only ten minutes, becoming *∼* 1 *µM* ; during this ten minutes GSH continues to decline at (almost, but not exactly) its previous rate. Then, at about *t* = 1.65 *hr*, H_2_O_2_ concentration jumps almost instantaneously by another order of magnitude, attaining very high values. This coincides with a “kink” in the plot of GSH concentration versus time, after which point GSH declines more slowly.

**Figure 3:**
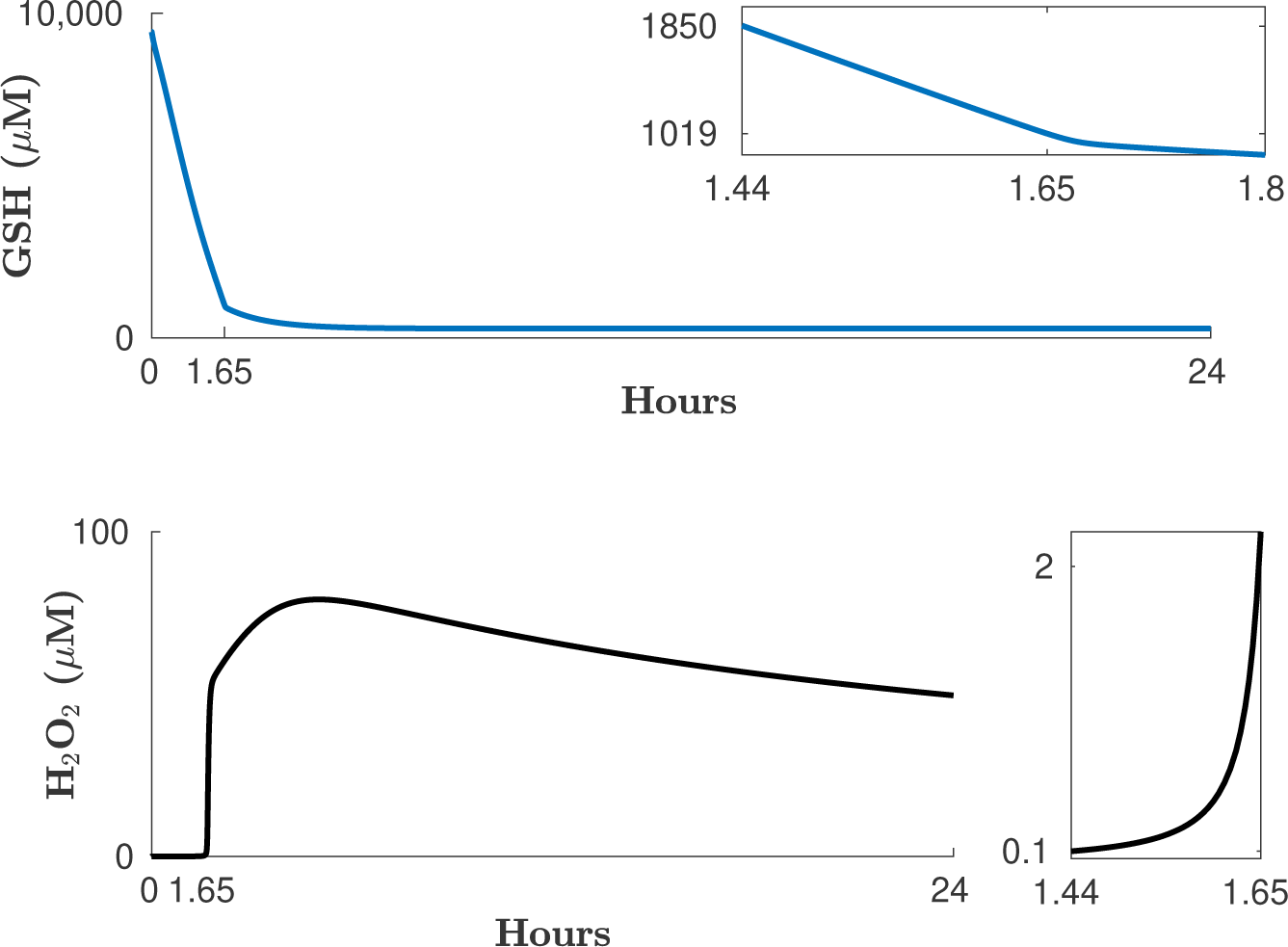
Simulation of Naranmandura experiments: *in vitro* exposure of rat hepatocytes to 7.5 *µM* DMA*^III^* in medium and resultant intracellular GSH and H_2_O_2_ concentrations, without NAC supplementation. Intracellular DMA*^III^* impacts the functioning of endoplasmic reticulum and increases the rate of H_2_O_2_ production by this organelle. This results in a very slow rise in intracellular H_2_O_2_ concentration during the first hour and a half of DMA*^III^* exposure, from the normal value of 0.01 *µM* to around 0.1 *µM*. While this is occurring, the above-normal flux through the glutathione peroxidase (GPX) reaction depletes GSH, despite the fact that it is being recycled from GSSG via glutathione reductase (GR) and synthesized *de novo* from cysteine taken up from the cellular medium (Dulbecco’s Modified Eagle Medium). Once GSH falls below a critical concentration, at around *t* = 1.5 *hr*, the H_2_O_2_ concentration rises much more rapidly (see inset of lower panel, covering the time interval from *t* = 1.44 *hr* to *t* = 1.65 *hr*), entering the range associated with apoptosis. For the 7.5 *µM* DMA*^III^* exposure level, the critical concentration of GSH is near 1,500 *µM* (*≈* 15% normal). Shortly after *t* = 1.65 *hr*, GSH concentration becomes too low for glutathione peroxidase to contribute significantly to the removal of H_2_O_2_. At this point catalase is the only effective means of removing H_2_O_2_, and H_2_O_2_ concentration rises dramatically, due to the low affinity of catalase for H_2_O_2_. Without the action of catalase, simulations indicate that H_2_O_2_ would attain values causing necrosis (*≥* 500 *µM* [52]); hence, catalase enables apoptotic rather than necrotic cell death during incidents of extreme oxidative stress.

I explain these results as follows. An examination of the flux expression for glutathione peroxidase shows that the term featuring GSH is squared, reflecting the fact that two molecules of GSH are needed to neutralize one molecule H_2_O_2_. As a result, this term shrinks rapidly as cytoplasmic GSH comes within range of its *K_m_* value in the GPX reaction (1330 *µM*). For example, when cytoplasmic GSH is present at double the *K_m_*, the GSH-dependent term in the GPX flux expression is 0.44, as opposed to 0.77 at *t* = 0 *hr* when GSH is at its normal physiological concentration of about 9,500 *µM*. GSH concentration eventually becomes low enough to cause a significant mismatch between the ability of GPX to remove H_2_O_2_ and the rate at which H_2_O_2_ is produced. This drives a rapid rise in H_2_O_2_ concentration, to values for which the H_2_O_2_-dependent term in the GPX flux expression nearly compensates for the now-significantly-diminished GSH-dependent term, and GSH declines at nearly the previous rate. During this period (*t* = 1.44 *hr* to *t* = 1.65 *hr*) H_2_O_2_ concentration enters the range associated with apoptosis. In the jargon of dynamical systems, the time interval between about *t* = 1.44 *hr* and *t* = 1.65 *hr* comprises a “boundary layer,” a region of space or time over which a dependent variable undergoes rapid change. This boundary layer terminates in an even more dramatic second boundary layer, when GSH falls so low that rising H_2_O_2_ concentration cannot preserve the flux through GPX: the H_2_O_2_-dependent term in the flux expression for GPX has a saturating (Michaelis-Menten) shape, with *K_m_* for H_2_O_2_ of 6.8 *µM* ; once H_2_O_2_ concentration increases to *∼* 1 *µM* this term rapidly saturates. With GSH too low for GPX to make a nontrivial contribution to H_2_O_2_ removal, H_2_O_2_ concentration rises extremely rapidly, attaining the very high values necessary for the significant removal of H_2_O_2_ by catalase (*K_m_ ≈* 30, 000 *µM*).

#### 3.2.3. Insights into how NAC treatment raises by an order of magnitude the extracellular DMA^III^ concentration threshold for appreciable cell death by the 24-hr mark: intra-cellular GSH has attained supraphysiological concentration, and excess cysteine is still present in medium, by the time DMA^III^ exposure begins

In contrast to the non-NAC-treated (control) hepatocytes, for whom exposure to 7.5 *µM* DMA*^III^* in medium killed about 70% of cells by the 24-hour mark, nearly no cell death was observed in the NAC-treated hepatocytes for this exposure level, dramatically demonstrating the power of NAC to rescue these cells. Fig. 4 shows the time course of intracellular DMA*^III^* concentration computed under the model for the 7.5 *µM* DMA*^III^* exposure level, and the time courses of intracellular H_2_O_2_ concentration computed for the non-NAC-treated and the NAC-treated cells. As discussed previously, H_2_O_2_ concentration is predicted to enter the range associated with apoptosis in the non-NAC-treated hepatocytes, but the NAC-treated hepatocytes are predicted to experience only a trivial rise in intracellular H_2_O_2_ concentration. Fig. 6 presents the analogous simulation results for each of the exposure levels tested by Naranmandura et al. [43]. H_2_O_2_ concentration in the NAC-treated cells is predicted to enter the range associated with apoptosis only for the highest exposure level tested, 10 *µM* DMA*^III^* in medium (*Row 6, third column*), while H_2_O_2_ in the non-NAC-treated cells is predicted to enter this range when DMA*^III^* is present in medium at only one-tenth this concentration, 1 *µM* (*Row 2, second column*). See also Fig. 2, and compare the 10 *µM* -DMA*^III^* data point for the NAC-treated cells with the 1 *µM* -DMA*^III^* data point for the non-NAC-treated cells. Fig. 5 shows simulation results for the NAC-treated hepatocytes exposed to 7.5 *µM* DMA*^III^*; cf. Fig. 3. The presence of 2000 *µM* NAC in medium greatly stimulates, over the 24-hour period preceding DMA*^III^* exposure, the flux through the glutamate-cysteine ligase reaction (the rate-limiting step in GSH synthesis), so that by the time DMA*^III^* is introduced (at *t* = 0 *hr* in Fig. 5), GSH has attained supraphysiological concentration. Although its concentration rapidly decreases upon the introduction of DMA*^III^*, it remains above 5000 *µM* at all times during the 24-hour period examined, high enough to keep the H_2_O_2_ concentration in check. I emphasize that, under the model, at each point in time the intracellular DMA*^III^* concentration and, hence, the rate of H_2_O_2_ production in cells, is the same for the non-NAC-treated and NAC-treated cells.

**Figure 4:**
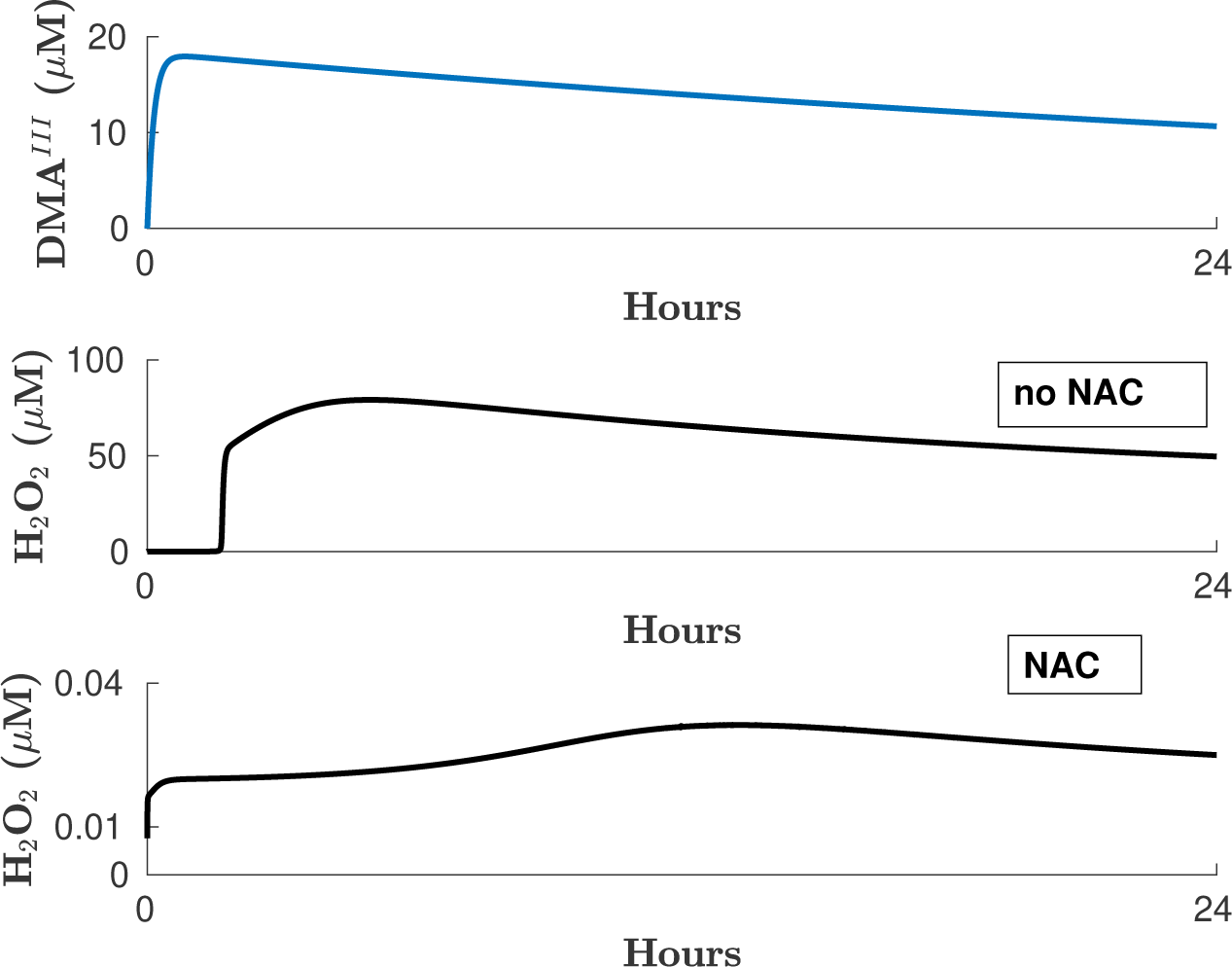
Simulation of Naranmandura experiments: *in vitro* exposure of rat hepatocytes to 7.5 *µM* DMA*^III^* in medium and resultant intracellular H_2_O_2_ concentration, with and without NAC supplementation. *Top*: Intracellular DMA*^III^* concentration as predicted by the *in vitro* PBPK model of DMA*^III^* and DMA*^V^* pharmacokinetics, Eqs. A.18-A.21. In the model, cells take up DMA*^III^* from medium and oxidize it to DMA*^V^* (not shown); DMA*^V^* is then exported to medium. *Middle*: Intracellular DMA*^III^* elevates above normal the rate of production of H_2_O_2_ by endoplasmic reticulum. Without N-acetyl cysteine (NAC, a cysteine source for the synthesis of GSH) supplementation, intracellular H_2_O_2_ attains concentrations causing (or possibly just associated with) apoptosis (*>* 1 *µM*). *Bottom*: When cells are cultured for 24 hours in medium supplemented with 2,000 *µM* NAC prior to DMA*^III^* exposure, this dramatic increase in H_2_O_2_ concentration is not seen.

**Figure 5:**
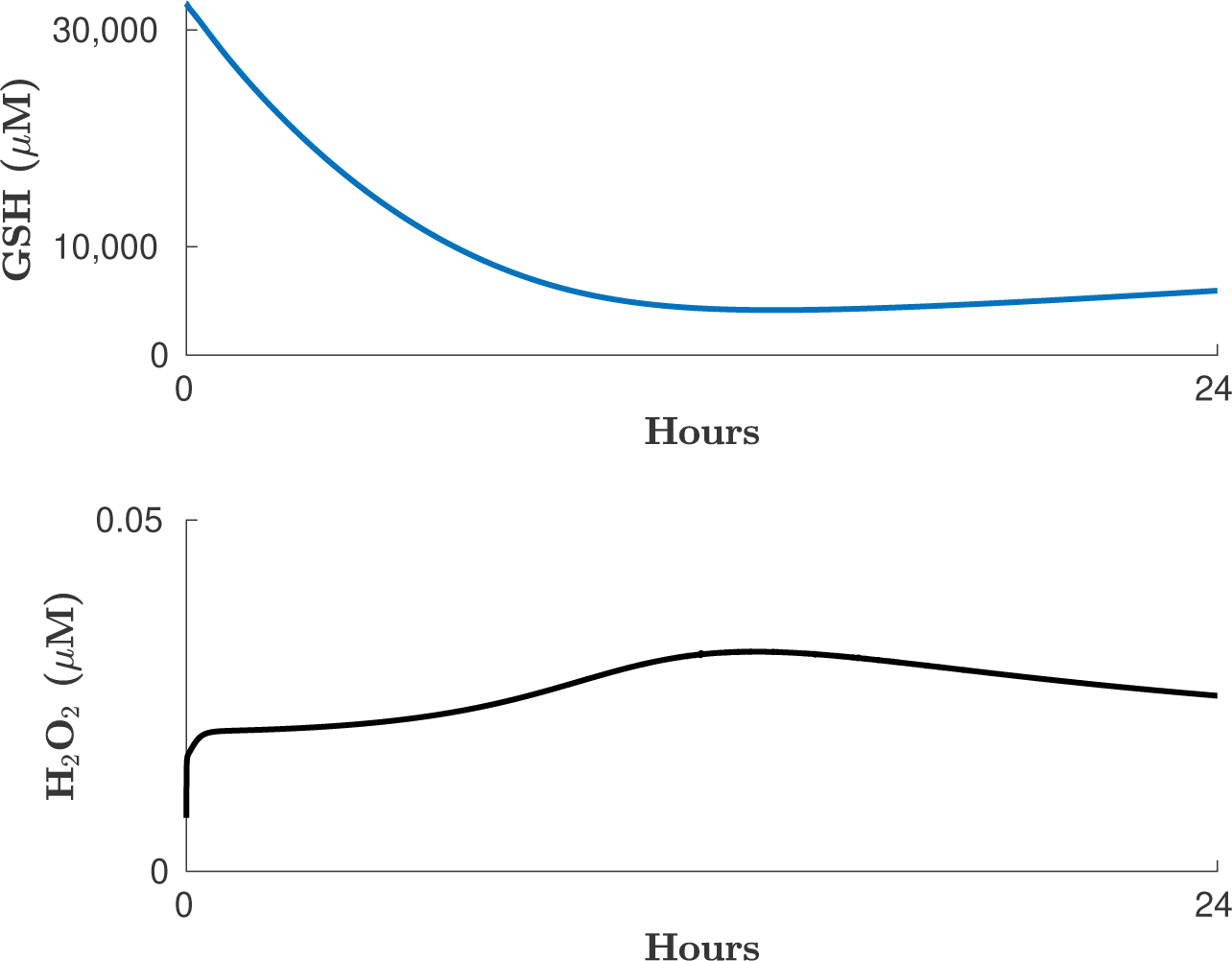
Simulation of Naranmandura experiments: *in vitro* exposure of rat hepatocytes to 7.5 *µM* **DMA***^III^* in medium and resultant intracellular GSH and H_2_O_2_ concentrations, with NAC supplementation. Cf. Fig. 3. *Top*: The NAC-supplemented hepatocytes are cultured in medium containing (in addition to the usual cysteine and cystine present in Dulbecco’s Modified Eagle Medium) 2000 *µM* NAC over the 24-hour period immediately preceding the addition of DMA*^III^* to medium. As a result, by the time DMA*^III^* is introduced (at *t* = 0 *hr*), they have attained supraphysiological intracellular GSH concentration; in addition, NAC is still present in medium. This situation confers dramatic protection against oxidative stress. The model predicts that GSH declines dramatically upon the introduction of DMA*^III^* but remains above 5000 *µM*, well above the critical concentration threshold below which the ability of glutathione peroxidase to remove H_2_O_2_ becomes compromised. *Bottom*: Intracellular H_2_O_2_ concentration is not predicted to rise significantly.

**Figure 6:**
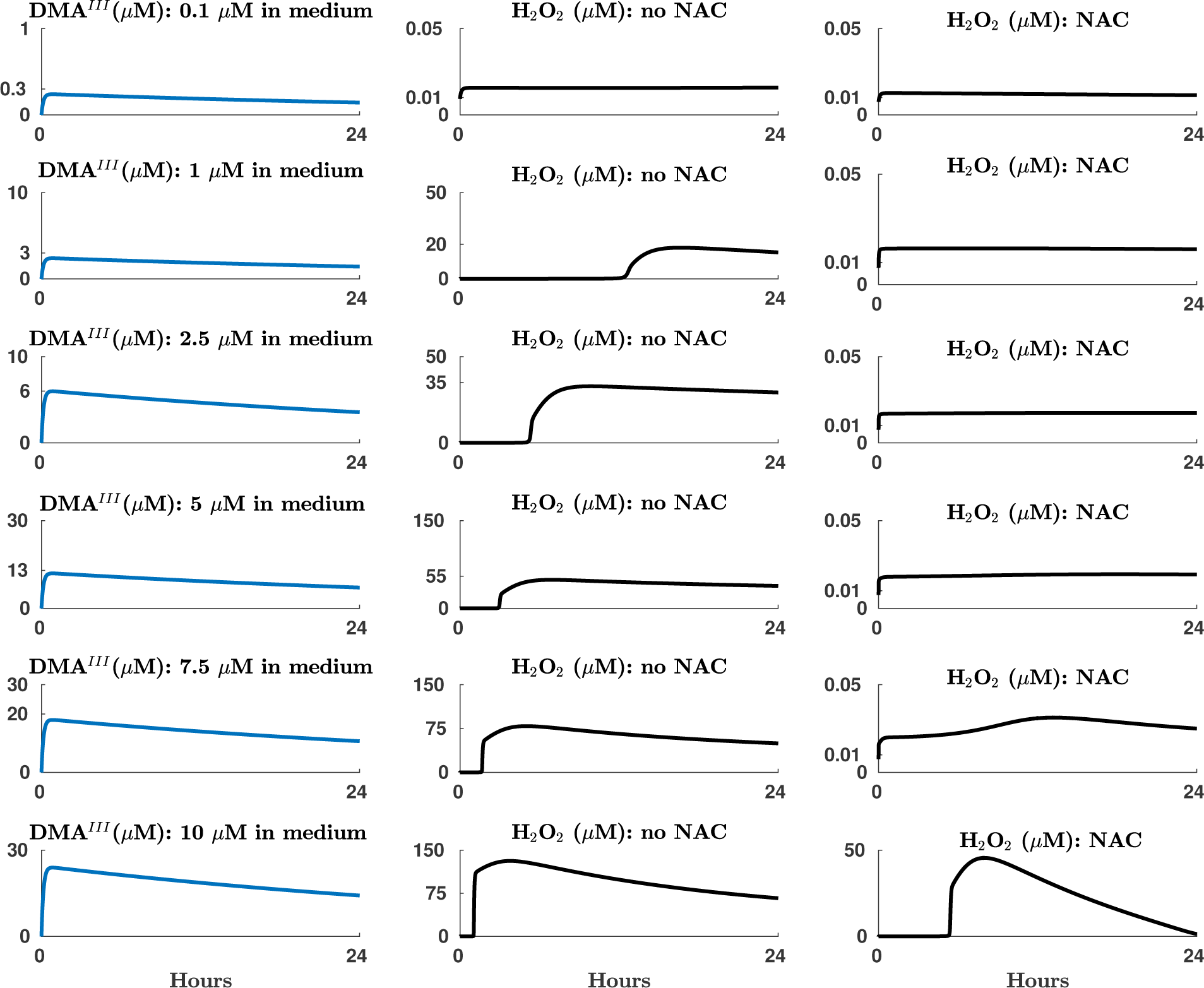
Simulation of Naranmandura experiments: *in vitro* exposure of rat hepatocytes to each tested concentration of DMA*^III^* in medium, and resultant intracellular DMA*^III^* and H_2_O_2_ concentrations, with and without NAC supplementation. Each row of this figure presents the model-predicted intracellular DMA*^III^* and H_2_O_2_ concentrations in rat hepatocytes exposed to a given concentration of DMA*^III^* in medium, as was done in the Naranmandura experiments [43]. The computational model predicts that exposure to 0.1 *µM* DMA*^III^* in medium does not result in an appreciable elevation in intracellular H_2_O_2_ above its normal physiological value of approximately 0.01 *µM* (*Row 1*). For all other tested exposure levels, H_2_O_2_ concentration eventually rises to values 3 to 4 orders of magnitude above normal in the non-NAC-treated cells, well within the range associated with apoptosis. The model predicts that NAC supplementation affords powerful protection against DMA*^III^* -induced oxidative stress: H_2_O_2_ concentration in the NAC-treated cells only enters the apoptotic range for the highest exposure level tested, 10 *µM* DMA*^III^* in medium (*Row 6*, third column). In contrast, this already occurs in the non-NAC-treated cells for the 1 *µM* DMA*^III^* exposure level. Note the existence of a time delay between DMA*^III^* reaching its peak concentration and H_2_O_2_ concentration entering the apoptotic range, in those cases where it eventually does; this is discussed in 3.2.4 in the text.

#### 3.2.4. Existence of a time delay between peak intracellular DMA^III^ concentration (which coincides, under the model, with the peak rate of H_2_O_2_ production) and H_2_O_2_ concentration entering the apoptotic range. Discovery of a GSH-mediated mechanism enabling cells to (1) distinguish normal, transient elevations in intracellular H_2_O_2_ concentration from pathological ones and (2) send cells experiencing the latter toward apoptosis

An inspection of Fig. 6 indicates that the model predicts a curious phenomenon: (1) for those cases in which H_2_O_2_ concentration enters the range associated with apoptosis during the 24-hour period considered, the time at which this occurs (and cell death commences) does not coincide with the time point at which intracellular DMA*^III^* concentration peaks, even though the latter is in fact when the rate of production of H_2_O_2_ is maximal, but instead comes later; (2) this “time delay” increases as exposure level decreases for all DMA*^III^* concentrations found to cause significant cell death within the 24-hour period examined. For example, for the case of non-NAC-treated cells exposed to 7.5 *µM* DMA*^III^*, the time delay is about an hour and a half, whereas for the 1 *µM* DMA*^III^* exposure level, it is about 12 hours. The delay is reflective of the “induction period” during which GSH is in decline but not yet critically depleted. The induction period is longer for lower DMA*^III^* exposure levels because they effect a smaller immediate elevation in H_2_O_2_ concentration over its normal value, so that, although it is high enough to eventually cause critical GSH depletion, this does not occur as rapidly as it does at the higher DMA*^III^* exposure levels. Very low DMA*^III^* exposure levels (e.g. 0.1 *µM*) can be indefinitely tolerated, because the increase they effect in the rate of H_2_O_2_ production is mild enough for the system to reach a new steady state, in which H_2_O_2_ concentration is very mildly elevated above its normal physiological value. I note that the function *f* ([DMA*^III^*]) is not sigmoidal (even trace intracellular DMA*^III^* is predicted to elevate the rate of H_2_O_2_ production in cytoplasm).

To summarize, the model predicts that a nontrivial elevation in the intracellular H_2_O_2_ concentration above its normal physiological value will eventually (if it persists) trigger apoptosis, with critical GSH depletion serving as the apoptotic trigger; smaller elevations in H_2_O_2_ concentration will be tolerated for a longer period of time, as they take longer to drive this critical depletion. This provides a GSH-based mechanism for cells to take into account both the intensity and the time persistence of an elevation in intracellular H_2_O_2_ concentration in “deciding” whether the cellular situation underlying the elevation is pathological (due to the presence of a xenobiotic or damage to organelles, etc.) and should end in apoptotis. Hence, the role/function of GSH within this mechanism is to become depleted. This mechanism helps to explain how intracellular H_2_O_2_ concentration is so tightly controlled (kept within such a narrow concentration range).

### 3.3. Even when no xenobiotic is present, extreme GSH depletion alone is predicted to trigger intracellular H_2_O_2_ concentration in the range associated with apoptosis

Simulation of Naranmandura and coworkers’ DMA*^III^* experiments enabled me to identify not only the function *f* but also the parameter *death* dictating cellular death rate as a function of intracellular H_2_O_2_ concentration. As mentioned earlier, whether intracellular H_2_O_2_ concentration in the range associated with apoptosis (*>* 1 *µM*) is a cause or merely a correlate of apoptosis is a matter of controversy; for simplicity, I assume here that the relationship is causal. Having identified the value of *death*, I returned to the case of an *in vitro* preparation of rat hepatocytes when no xenobiotic is present (no DMA*^III^*) to investigate the cellular consequences of extreme GSH depletion alone. Using the steady-state concentrations in Table 1 as the initial condition, I set *V_max_* for the glutamate-cysteine ligase reaction (the first step in glutathione synthesis) to zero in order to simulate a situation in which GSH synthesis is totally halted. Results are shown in Fig. 7. The model predicts that the plentiful store of GSH under normal physiological conditions (about 9,500 *µM*) and the ability of glutathione reductase to recycle GSSG back to GSH enables hepatocytes to control intracellular H_2_O_2_ concentration in the face of its production by endogenous processes (production by mitochondria and cytoplasmic organelles) for 12 hours after *de novo* GSH synthesis has been halted. This is even more impressive given that export of GSH and GSSG to medium is still taking place. After roughly 12 hours, H_2_O_2_ levels are predicted to enter the apoptotic range. Simulation results are consistent with various empirical findings, without having been contrived for that purpose: (1) Hepatocytes in which GSH has been depleted, such as by treatment with diethyl maleate (DEM), a chemical which conjugates with and removes GSH, can experience a depletion in GSH of 95% without an immediate loss of viability [49]. (2) Rapid cell death occurs when intracellular H_2_O_2_ concentration is of order 10 *µM* [59]. (3) The GSH:GSSG ratio (glutathione redox ratio), usually of order 100:1, decreases to 10:1 or even 1:1 during conditions of oxidative stress [12]. This serves as a validation of the model. I note that, for the rat hepatocytes exposed to 7.5 *µM* DMA*^III^* in medium (Fig. 3), intracellular H_2_O_2_ concentration entered the apoptotic range once GSH was only about 85% depleted (rather than more than 95% as here); the elevated rate of production of H_2_O_2_ in these DMA*^III^*-containing cells raised the GSH concentration threshold below which H_2_O_2_ concentration enters the range associated with apoptosis.

**Figure 7:**
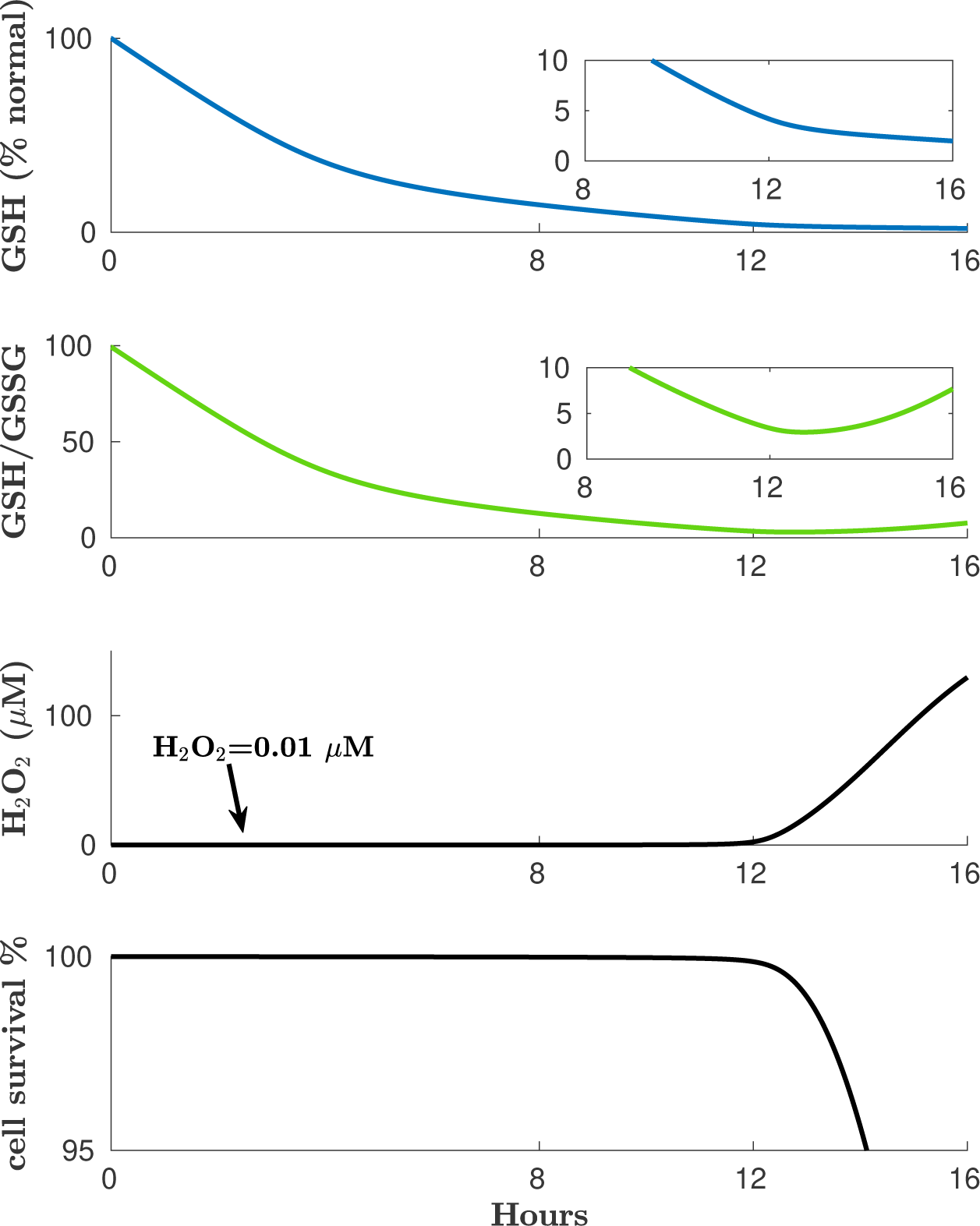
Model-predicted effects of halting *de novo* GSH synthesis in rat hepatocytes. Concentrations are total-cell averages (over mitochondrial and cytoplasmic compartments). *Top*: GSH concentration expressed as a percentage of the normal steady-state value. *Second from top*: Ratio of GSH to GSSG concentrations (called the “glutathione redox ratio”). *Third from top*: Intracellular H_2_O_2_ concentration. *Bottom*: Hepatocyte survival, where the parameter *death* in Eq. A.23 has the value given in Table A.4. Solutions are consistent with the fact that GSH in hepatocytes can be depleted by even 95% without an immediate loss of viability [49]. Solutions are also consistent with the fact that the GSH:GSSG ratio, usually *≈* 100:1, decreases to 10:1 or even 1:1 under conditions of oxidative stress [12]. The increase in this ratio after the 12-hour mark reflects the loss of GSSG via its efflux from hepatocytes, rather than a genuine recovery of reduced conditions.

### 3.4. in vivo computational model of endogenous H_2_O_2_ production and removal in mouse liver (no xenobiotic present)

As a preliminary to creating a PBPK/PD model for dosing of the mouse with DMA*^III^* or DMA*^V^*, I adapted the *in vitro* computational model of endogenous H_2_O_2_ production and removal in rat hepatocytes to the case of mouse liver *in vivo*. Table 2 shows the concentrations at steady state for this model. Concentrations in blood plasma and liver tissue agree well with values in the published literature.

**Table 2:** Steady-state concentrations of biochemical substrates computed for the *in vivo* computational model of endogenous H_2_O_2_ production and removal in mouse liver (no xenobiotic present).

| Model concentration ($\mu M$) | Normal physiological range/comment | Source |
| --- | --- | --- |
| **Intracellular** |  |  |
| $H_2O_{2,cyt}=0.01$ | 0.01-0.1 | [8] |
| $H_2O_{2,mito}=0.005$ | 0.005 | [8] |
| $GSH_{cyt}=10030$ | 9886-13043 | [45] |
| $GSH_{mito}=9989$ | similar to cytoplasmic value | [51] |
| $GSSG_{cyt}=100$ ($\approx 2\%$ total GSH equivalents) | 0.58-1.3% (NEM method, whole liver) | [38] |
|  | 1.5-8.2% (2-VP method, whole liver) | [38] |
| $GSSG_{mito}=99$ ($\approx 2\%$ total GSH equivalents) | 1.25-2.44% (mitochondria of mouse) | [48] |
| γ-GC=9.86 | | |
| Cys=120 | 96-136 | [68] |
| **Plasma in liver capillary bed** |  |  |
| GSH=122 |  |  |
| Cys=0.07 |  |  |
| **Blood plasma** |  |  |
| GSH=100 | 56-138 | [68] |
| Cys=24 | 8-24 (10-20% total plasma cysteine equivalents) | [68],[53] |
| Cyss=50 | 34-52 (80-90% total plasma cysteine equivalents) | [68],[53] |

### 3.5. in vivo PBPK/PD model predicting H_2_O_2_ concentration in mouse liver and hepatocyte death after oral or intravenous dosing of mouse with DMA^III^ or DMA^V^

I conjoined the *in vivo* computational model of endogenous H_2_O_2_ production and removal in mouse liver to my and coworkers’ 2019 PBPK model for oral or intravenous dosing of mouse with DMA*^III^* or DMA*^V^* [7]. The point of juncture between the two models is the function *f* ([DMA*^III^*]) identified during the construction of the *in vitro* PBPK/PD model. Here I am assuming that mouse and rat hepatocytes have the same response to arsenicals; this work serves as a proof of concept which can be made more accurate once *in vitro* and *in vivo* data sets become available for a single species.

#### 3.5.1. Simulation of oral dosing of mouse with its LD50 for DMA^V^

I used the *in vivo* mouse PBPK/PD model to predict the consequences for mouse liver of oral dosing of mouse with its LD50 for DMA*^V^* (650 mg As/kg) [54]. Results are shown in Fig. 8. Bilinsky et al. predicted [7] that orally administered DMA*^V^* is extensively reduced in the gut of mouse to DMA*^III^*, so that both valence forms enter liver and systemic circulation; this has now been shown experimentally [67]. The top panel of Fig. 8 shows the predicted intracellular DMA*^III^* and DMA*^V^* concentrations resulting from this dose. Recall that it is only the trivalent form (DMA*^III^*) that is responsible for the generation of ROS in hepatocytes; DMA*^V^* is not modeled as impacting ROS production.

**Figure 8:**
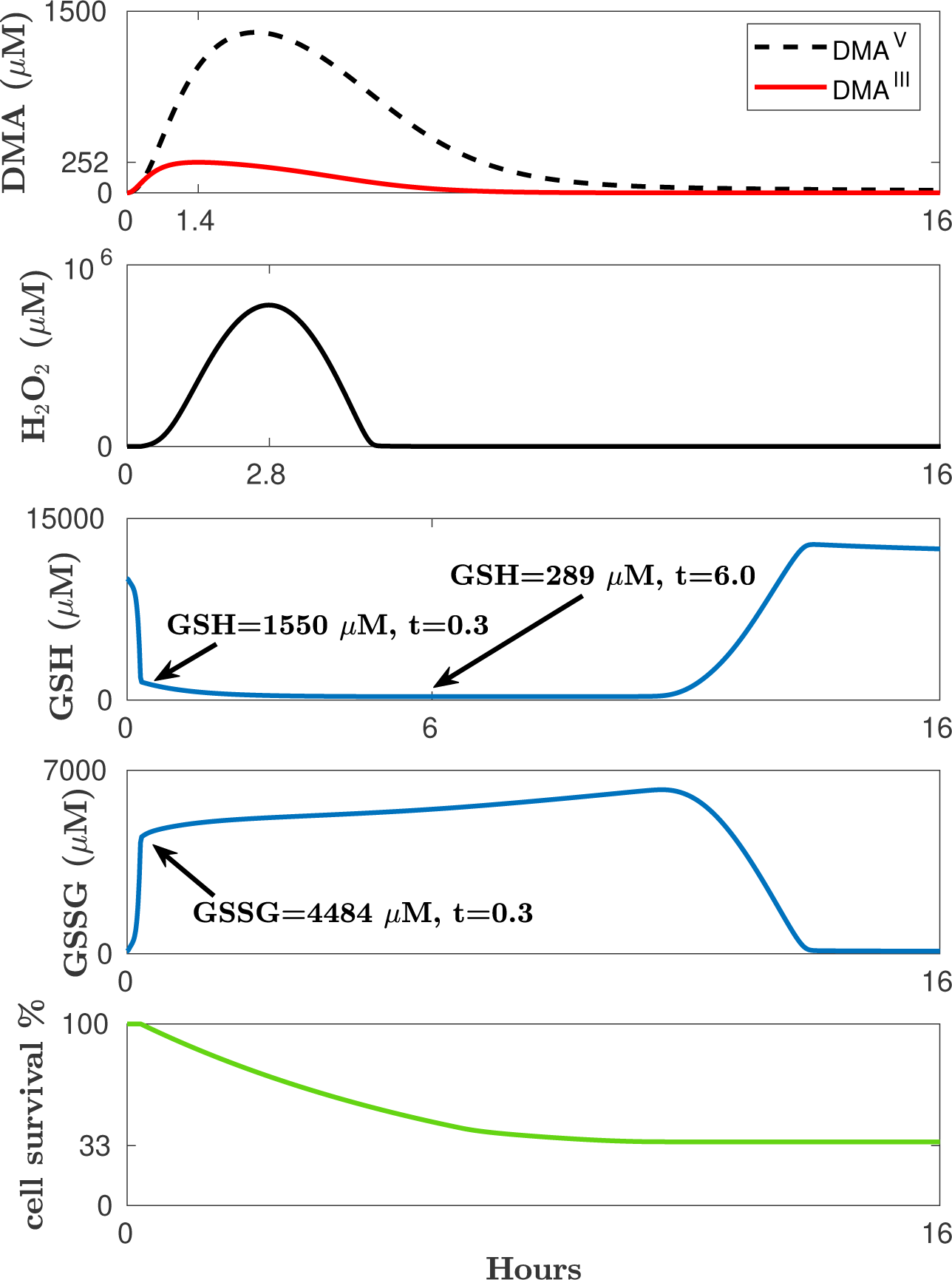
Model-predicted consequences for mouse liver of oral dosing of mouse with 650 mg As/kg DMA*^V^* (the oral LD50 [54]). *First panel* : Predicted intracellular DMA*^III^* and DMA*^V^* concentrations in liver. Orally administered DMA*^V^* undergoes extensive reduction in intestine of mouse to DMA*^III^* [7, 67]; hence, both valence forms enter liver. *Second panel* : Predicted intracellular H_2_O_2_ concentration. Although this is unphysiologically high, for reasons discussed in the text (see 3.5.1), this simulation provides useful insights as it illustrates the most extreme case of xenobiotic-induced oxidative stress. Note that the peak in H_2_O_2_ concentration takes twice as long to arrive as the peak in DMA*^III^* concentration (“delay effect”), even though the latter is when the rate of intracellular H_2_O_2_ production is greatest. *Third and fourth panels*: The model predicts that intracellular GSH holdings will be shifted almost entirely into oxidized form (GSSG) during this intense oxidative assault, and that GSH will become extremely depleted despite its being intensively synthesized *de novo* at all times. GSSG is effectively trapped inside hepatocytes during the oxidative assault, due to its low *K_m_* for export (400 *µ*M [3]); this minimizes the need for *de novo* GSH synthesis to restore normal GSH concentration once the assault ends. *Fifth panel* : Predicted percentage of living hepatocytes.

As GSH concentration (third panel) falls and DMA*^III^* concentration rises, the H_2_O_2_ concentration (second panel) increases, peaking near 1 *M*. It is unlikely that this is physiologically realistic; I show this result as an illustration of the most extreme scenario of xenobiotic-induced oxidative stress. This predicted H_2_O_2_ concentration stems from extrapolation of the function *f* ([DMA*^III^*]), containing a super-quadratic term, to intracellular DMA*^III^* concentrations far above the range attained in Naranmandura and coworkers’ experiments (see first column of Fig. 6), over which *f* was obtained. This demonstrates the need to collect more *in vitro* data for DMA*^III^*, using higher DMA*^III^* exposures and exposure periods less than 24 hours. I note also that mouse hepatocytes can be expected to respond at least somewhat differently to DMA*^III^* than do rat hepatocytes and that ideally *in vitro* and *in vivo* data sets for a single species should be used. Nevertheless, this simulation provides useful mechanistic insights, discussed next.

Fig. 8 shows that DMA*^III^* peaks at *t* = 1.4 *hr* after dosing but, due to ongoing the decline in GSH, H_2_O_2_ concentration does not peak until *t* = 2.8 *hr*. GSH is still in decline at *t* = 2.8 *hr* but DMA*^III^* is now low enough for the flux through GPX to clear H_2_O_2_ faster than it is produced, and H_2_O_2_ concentration begins to decline. The fifth panel shows that approximately 2/3 of hepatocytes have died by the time the DMA*^III^* and DMA*^V^* are eliminated. Studies of hepatectomy in mouse have determined that the liver can regenerate even after 70% removal [26]. The death of 2/3 of hepatocytes in response to an LD50 dose is a result that I achieved, for illustrative purposes, by assuming that the per-cell death rate of hepatocytes, modeled as increasing linearly with H_2_O_2_ concentration, attains a maximal value once [H_2_O_2_]=150 *µM* (see 4). This result serves as a proof of concept; in future work, utilizing *in vitro* and *in vivo* data sets for a single species and for which the exposure concentrations/doses investigated translate into similar intracellular DMA*^III^* concentrations, the H_2_O_2_ concentration ceiling on cellular death rate should either be discarded, or set to a value chosen according to independent considerations. I remark that in the *in vitro* and *in vivo* computational models presented in this paper, hepatocyte regeneration (replication) is neglected, due to the short time scale considered, and due to a departure from cellular conditions that are conducive to cell replication, caused by the presence of large amounts of DMA*^III^* (e.g. interference with normal functioning of the endoplasmic reticulum).

Panels three and four of Fig. 8 show that the immediate consequence of this oral dose of DMA*^V^* is a conversion of nearly the entire liver GSH store to the oxidized form GSSG (recall that two molecules of GSH are consumed in producing one GSSG). GSSG then continues to rise more slowly, at rate reflective of the rate of *de novo* GSH synthesis (which is immediately consumed in reaction with H_2_O_2_), until about the 12-hour mark. At this time, the concentration of H_2_O_2_ is finally low enough for the flux through GPX to cease to dominate the flux through GR, and GSSG is then re-converted to GSH. Although GSSG can be exported from hepatocytes via a transporter, its *K_m_*for GSSG is only 400 *µM* ; hence, export is rapidly saturated as intracellular GSSG rises. I speculate that the low *K_m_*for GSSG export is an evolved mechanism to trap GSSG inside hepatocytes during incidents of extreme oxidative stress, minimzing the amount of *de novo* synthesis needed to restore GSH concentration to its normal 10 *µM* once the oxidative assault ends. In fact, panel four of Fig. 8 shows that intracellular GSH will reach concentration about 30% higher than normal, reflective of the original GSH store plus the GSH molecules made and consumed during the oxidative insult. GSH efflux then slowly restores liver concentration to its normal physiological value. I emphasize that even when the computed intracellular GSH is extremely depleted (e.g., at the 6-hour mark, when it is predicted to be present at only 300 *µM*) GSH is being intensively synthesized *de novo* at rate even exceeding normal, due to the lifting of product inhibition on GCL, and also regenerated from GSSG via GR. Hence, simulations predict that GSH can become extremely depleted during incidents of extreme oxidative stress, with kinetic consequences for the rate at which H_2_O_2_ can be removed.

#### 3.5.2. Tremendous ability of undepleted GSH to control intracellular H_2_O_2_ concentration in the face of an elevated rate of production

Fig. 9 shows what happens if I rerun the oral LD50 simulation, this time holding GSH concentration constant in mitochondria and cytoplasm but making no other changes. H_2_O_2_ concentration at its highest is now a few *µM*, versus 100,000 times greater as in Fig. 8; this demonstrates the incredible ability of GSH, when sustained at its normal concentration, to remove H_2_O_2_. Hence, it is not a limitation of the capacity of the GSH-based antioxidant system to utilize GSH to remove H_2_O_2_ that underlies H_2_O_2_ concentration entering the apoptotic or necrotic range when an oxidative-stress-producing xenobiotic such as DMA*^III^* is present, but rather, a limitation in the ability of the hepatocyte to generate GSH at a rate matching its rate of consumption under these circumstances, either by synthesizing it *de novo* from glutamate, cysteine, and glycine, or by recyling GSSG to GSH using the enzyme glutathione reductase (GR).

**Figure 9:**
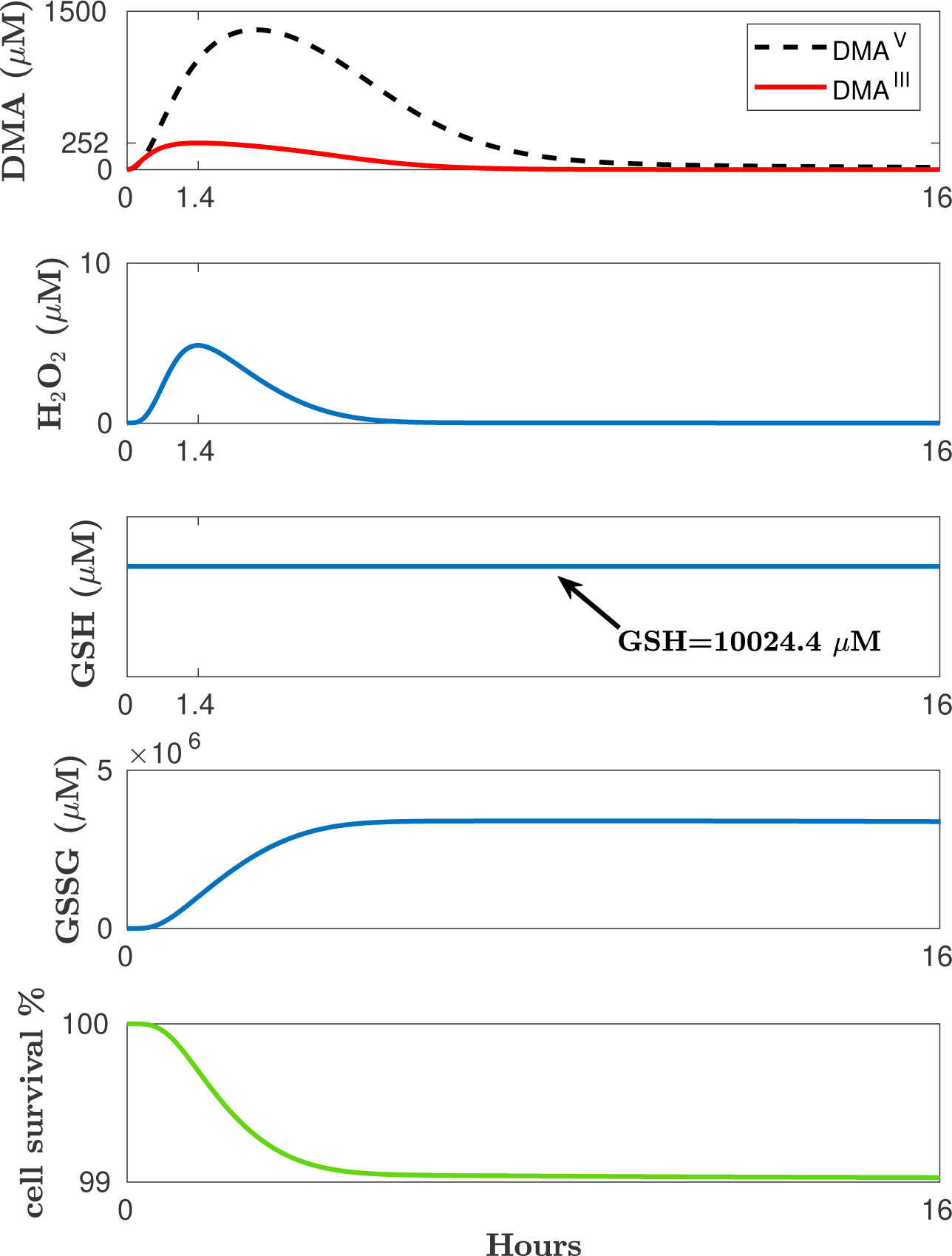
Model-predicted consequences for mouse liver of oral dosing of mouse with its LD50 for DMA*^V^* when cytoplasmic and mitochondrial GSH are numerically held constant; cf. Fig. 8. The system solved here is identical to that solved in Fig. 8, except that the mitochondrial and cytoplasmic GSH concentrations are numerically held constant at their normal physiological values. The peak intracellular H_2_O_2_ concentration is 100,000 times smaller than was predicted in Fig. 8, and coincides with the peak in DMA*^III^* concentration (no delay effect, the mechanism of which is decreasing GSH concentration). For a detailed discussion of these results, see 3.5.2.

#### 3.5.3. Existence of a time delay between peak intracellular DMA^III^ concentration and peak intracellular H_2_O_2_ concentration in mouse liver after oral dosing with DMA^V^ ; comparison of in vivo and in vitro cases

Panels one and two of Fig. 8 show that the peak in intracellular H_2_O_2_ concentration is predicted to take twice as long to arrive as the peak in intracellular DMA*^III^* concentration, despite that fact that the latter is when the rate of production of H_2_O_2_ in cells is greatest. This time delay effect was also observed for simulations of Naranmandura’s *in vitro* experiments with rat hepatocytes (see Fig. 6). Note that much lower intracellular DMA*^III^* concentrations are predicted for those experiments than for oral dosing with DMA*^V^*, but that the DMA*^III^* is predicted to be present for a much longer time. In *in vivo* exposure scenarios, DMA*^III^* and DMA*^V^* rapidly enter liver via the portal vein while excreted DMA*^V^* is carried away by the bloodstream, to venous drainage. Some second-pass metabolism occurs but arsenical exposure is not nearly so prolonged as in the “stagnant” situation of hepatocytes exposed to DMA*^III^* in an *in vitro* preparation. The latter experience a less pronounced, but much longer in duration, elevation in the rate of H_2_O_2_ production over baseline, resulting in a longer time to critically deplete GSH; in fact, it takes about 12 hours in the case of the cells exposed to 1 *µM* DMA*^III^*.

#### 3.5.4. Comparison of the in vivo PBPK/PD mouse model presented here with the earlier one in Bilinsky et al. [7], which did not feature depletion of the cofactor GSH

In [7], to demonstrate how our PBPK model of oral or intravenous dosing of mouse with DMA*^III^* or DMA*^V^* could be augmented to describe the pharmacodynamic action of DMA*^III^* in liver, thus generating a PBPK/PD model, I conjoined to it two additional equations: (1) a very simplistic model of endogenous H_2_O_2_ production and removal and of the generation of oxidative stress by DMA*^III^* and (2) an equation modeling cell death, identical to the one used in the current model (except for the value of the parameter *death*). The equation for H_2_O_2_ is given below.

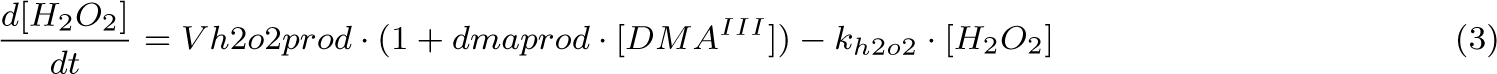

No distinction is made between mitochondrial and cytoplasmic compartments in that model, and *V h*2*o*2*prod* in the above equation refers to the whole-liver rate of endogenous production of H_2_O_2_. H_2_O_2_ removal is simply modeled as being linear in H_2_O_2_ concentration, with constant coefficient *k_h_*_2*o*2_, chosen to agree with steady-state concentrations. This modeling choice has the effect of greatly overestimating the cell’s ability to remove H_2_O_2_ during incidents of oxidative stress, since depletion of the cofactor GSH (which occurs during these incidents) is not modeled. Another difference is that a simple linear relationship is assumed in that model between intracellular DMA*^III^* concentration and the rate of production of H_2_O_2_; here, it was found, by fitting to Naranmandura and coworkers’s data sets, that the function *f* ([DMA*^III^*]) dictating this relationship is highly nonlinear. Finally, and very importantly, the range of intracellular H_2_O_2_ concentration associated with cell death was greatly overestimated in [7]: there, it was assumed to be what is required of *extracellular* H_2_O_2_ concentrations. Sies states that the ratio of extracellular H_2_O_2_ concentration to intracellular H_2_O_2_ concentration in cell preparations exposed to H_2_O_2_ is about 100:1 and that apoptotic cell death is seen for intracellular H_2_O_2_ concentration on order of 1-10 *µM* [59]. The current model includes this important correction.

The primary purpose of our 2019 paper was to present a pharmacokinetic model for DMA*^III^* and DMA*^V^* in mouse; the pharmacodynamic extension was just a proof of concept. The current PBPK/PD model is far more realistic, in terms of model structure, parameter values, and indeed behavior. In [7] the value, obtained from optimization, for the parameter *dmaprod* appearing in Eq. 3 is 1500 *µM ^−^*^1^, implying that an intracellular DMA*^III^* concentration of just 1 *µM* causes the rate of production of H_2_O_2_ in cytoplasm to be 1501 times the endogenous rate, which is clearly not realistic. Under the current model, this factor is approximately 3, since *f* (1 *µM* DMA*^III^*) *≈* 2. As another example, in [7] I simulated the impact on mouse liver of a 60 mg As/kg oral dose of DMA*^V^*, predicting that this dose kills 1/3 of liver hepatocytes. This was the higher of two oral doses administered by Hughes et al. in their pharmacokinetic studies of DMA*^V^* in mouse [28]; their data sets were used to construct the pharmacokinetic model in [7]. Under the current model, only trivial cell death is predicted, a much more realistic result given that this dose is less than 1/10 the LD50 dose of 650 mg As/kg [54].

## 4. Conclusions

The first deliverable of this paper is a relatively simple mathematical model of endogenous H_2_O_2_ production and removal in hepatocytes, of use in the construction of pharmacodynamic models for xenobiotics known to promote oxidative stress in liver. At the time of writing, it is unique among models of intracellular H_2_O_2_ metabolism found in the published literature in its suitability for this purpose, due to its ability to describe, relatively accurately, H_2_O_2_ dynamics in hepatocytes even during incidents of extreme oxidative stress, such as occur in the toxicological setting. The model is a set of ordinary differential equations (ODE). Separate compartments are used for mitochondria and cytoplasm. Model simulations predict the intracellular concentrations in hepatocytes of biochemical reactants relevant to H_2_O_2_ metabolism and the fraction of hepatocytes still living (relative to the starting number) at time *t*. Versions of the model are presented for the cases of rat hepatocytes *in vitro* and mouse liver *in vivo*. The model features the removal of H_2_O_2_ by the enzyme catalase (CAT), which requires no cofactor, and by the enzyme glutathione peroxidase (GPX), which requires the cofactor GSH (reduced form of glutathione). Given the importance of this cofactor, pains are taken to accurately model its dynamics. This is achieved by subsuming into the model a minimal version of the pioneering model of liver glutathione metabolism due to Reed et al. [50]. The modeling of H_2_O_2_ dynamics is heavily informed by work due to Sies and coworkers. From their papers, I (1) infer the rates of H_2_O_2_ production in mitochondria and in cytoplasm (by extramitochondrial cytoplasmic organelles, including endoplasmic reticulum) and (2) locate values for hydrogen peroxide’s normal physiological concentrations in mitochondria and in cytoplasm, as well as (3) the intracellular concentration range associated with apoptosis. The model is constructed to yield steady-state concentrations of featured biochemical reactants that agree with normal physiological values found in the published literature; e.g., GSH is present at about 9500 *µM* in rat hepatocytes (10000 *µM* in mouse hepatocytes) and the redox ratio of GSH:GSSG is about 100:1 under normal circumstances. Using the model to simulate a situation in which GSH synthesis is halted yields results consistent with the experimental findings that rat hepatocytes can experience a 95% depletion of GSH without an immediate loss of viability [49], and that during extreme oxidative stress the GSH:GSSG ratio becomes of order 10:1 or even 1:1 [12]. I note that the model was not constructed to yield these agreements, hence this serves as a validation.

Reed and coworkers’ 2008 model did not feature H_2_O_2_ dynamics as this was not their main priority, and H_2_O_2_ concentration was treated as a constant. Their model predicted that an above-normal H_2_O_2_ concentration drives a depletion of GSH via enhanced flux through GPX. A major result of the model presented here is that the converse is also true: depletion of GSH to a critically low level compromises the ability of GPX to remove H_2_O_2_, triggering a rapid rise in H_2_O_2_ concentration to levels associated with apoptosis (*>* 1 *µM* [59]). Fig. 7 shows that, for rat hepatocytes, this critical GSH concentration is less than 5% of the normal concentration if no xenobiotic promoting oxidative stress is present. If GSH falls below this, even the endogenous (normal physiological) production of H_2_O_2_ by mitochondria and by extramitochondrial cytoplasmic organelles is enough to cause an intracellular H_2_O_2_ concentration in the range associated with apoptosis. I note that when ROS-promoting xenobiotics are present, the critical GSH concentration threshold is higher. Fig. 3 shows that it is about 15% normal for rat hepatocytes exposed to 7.5 *µM* DMA*^III^* in medium, as the presence of intracellular DMA*^III^* causes the rate of production of H_2_O_2_ to exceed its normal physiological value.

The second deliverable is the demonstration of a generic procedure for constructing an *in vivo* PBPK/PD model for a xenobiotic known to promote oxidative stress in liver, using *in vitro* cell mortality data sets: (1) First, an *in vitro* PBPK model for the xenobiotic is conjoined to the *in vitro* version of the computational model of endogenous H_2_O_2_ metabolism in hepatocytes presented here, with juncture point a function *f* which must be identified, relating the rate of intracellular H_2_O_2_ production to intracellular xenobiotic concentration. *f* will appear either in the equation governing H_2_O_2_ concentration in mitochondria (if that is the target organelle, as is the case for MMA*^III^*) or, as in this case, for DMA*^III^*, in the equation governing H_2_O_2_ concentration in the cytoplasm. (2) The function *f* is identified by fitting the model to *in vitro* cell mortality data sets of the type collected by Naranmandura et al., reporting on the degree of cell death after a set number of hours’ exposure of cells to various concentrations of the xenobiotic in medium, with or without prior culturing with NAC. These data are extremely helpful because they enable us to use the quantitative information about reaction fluxes stored within the computational model of endogenous H_2_O_2_ production and removal in hepatocytes to infer the additional oxidative strain (rate of H_2_O_2_ production in excess of the endogenous value) caused by the xenobiotic. Once *f* has been identified, one obtains an *in vitro* PBPK/PD model. (3) One can now construct an *in vivo* PBPK/PD model for the xenobiotic: an *in vivo* (i.e., liver of the living animal) PBPK model for the xenobiotic is conjoined to the *in vivo* version of the computational model of endogenous H_2_O_2_ metabolism in hepatocytes, with contact point the function *f*.

Interesting phenomena from a dynamical systems perspective are observed: (1) Boundary layers: model simulations show time intervals during which the H_2_O_2_ concentration rises by an order of magnitude within an extremely short time span (known as a “boundary layer” in the parlance of dynamical systems). For example, Fig. 3 features two boundary layers: the first occurs when GSH concentration becomes kinetically rate-limiting for the removal of H_2_O_2_ via glutathione peroxidase, and H_2_O_2_ concentration, which has been maintained below 0.1 *µM* for the first hour and half, rises to *∼* 1 *µM* within about ten minutes. This first boundary layer ends when GSH is so depleted that GPX can no longer contribute significantly to the removal of H_2_O_2_, at which time H_2_O_2_ concentration rises nearly instaneously, in a second boundary layer, to concentrations in the range 10-100 *µM*, high enough to get the attention of catalase. The very high concentrations H_2_O_2_ settles into after the second boundary layer stem from the fact that catalase, now solely responsible for the effective removal of H_2_O_2_, has much lower affinity for H_2_O_2_ than does GPX: *K_m_ ∼* 10, 000 *µM* versus *∼* 1 *µM*. That extreme GSH depletion is the immediate cause of H_2_O_2_ concentration entering the apoptotic range underlies a second interesting phenomenon, described now. (2) Time delay phenomenon: as a general rule, there is a time delay between the arrival of the intracellular DMA*^III^* peak (at which time the rate of H_2_O_2_ production is maximal) and the arrival of the H_2_O_2_ peak, except in simulations where GSH is held constant (in which case they coincide). GSH is normally present at about 10,000 *µM* and is at all times being recycled via GR and synthesized *de novo*; hence, it may take hours for the flux through GPX (which is higher than normal because of the mild immediate elevation in H_2_O_2_ effected by the presence of DMA*^III^*) to drive GSH below the critical concentration for triggering a rise in H_2_O_2_ to concentrations associated with apoptosis. This delay is longer for smaller but eventually-lethal (that is, eventually causing measurable cell death within the 24-hour period examined) concentrations of DMA*^III^*, as the depletion of GSH occurs more slowly, due to smaller H_2_O_2_ concentrations driving the flux through GPX during the “induction period.” For the simulation of rat hepatocytes exposed *in vitro* to 1 *µM* DMA*^III^* in medium, this time delay is predicted to be about 12 hours.

The most important results of this work from a cellular physiology perspective are as follows. (1) The delay effect reflects a GSH-based mechanism for taking into account the time persistence of a mildly elevated H_2_O_2_ concentration (e.g. 0.1 *µM*, which, when transient, serves various normal physiological purposes [59] but is pathological as a baseline) in “deciding” whether the cellular situation is pathological and should terminate in apoptosis. This mechanism essentially applies a “low-pass filter” to cellular H_2_O_2_ concentration: transients are disregarded while a persistent elevation leads to apoptosis. (2) Rodrigo Franco and coworkers have found that cells executing apoptosis in response to Fas ligand binding actively extrude GSH to effect its depletion, that this depletion precedes ROS accumulation, that GSH depletion regulates the cellular death machinery through processes not involving ROS, and that GSH depletion is necessary for apoptosis [21, 22]. They speculate that extreme GSH depletion may be the central ingredient of the apoptotic mechanism, with elevated ROS simply indicating this condition. Simulations are consistent with this speculation: under the model, intracellular H_2_O_2_ concentration in the range associated with apoptosis (*>* 1 *µM*) implies that GSH is extremely low, hence automatically guaranteeing the GSH depletion criterion for apoptosis. Conversely, extreme GSH depletion immediately triggers H_2_O_2_ concentration in this range. Hence, under the model, GSH depletion and apoptotic or necrotic H_2_O_2_ concentration co-occur. I emphasize that although simulations are consistent with the hypothesis that H_2_O_2_ concentration *>* 1 *µM* is purely an indicator of an apoptotic cellular milieu, they do not require it, and it may be that elevated ROS do play a direct role in the apoptotic mechanism. (3) Simulations of oral dosing of mouse with DMA*^V^* indicate that, during incidents of extreme oxidative stress in liver, in which large amounts of GSSG are produced, the low *K_m_* value of the hepatic GSSG exporter serves to trap oxidized-form glutathione inside hepatocytes. I speculate that this is an evolved mechanism which preserves the glutathione pool during such incidents, minimizing the need for *de novo* GSH synthesis to restore normal GSH levels once the oxidative insult ends. (4) In this first model, NADPH, the cofactor needed by glutathione reductase to recycle GSSG back to GSH, is treated as constant. Realistically it can be expected to become depleted. The current model’s probable overestimate of the ability of the cell to maintain glutathione in reduced form via recycling proves the point even more dramatically that GSH can become critically depleted during the scenarios of extreme oxidative stress expected to occur during exposure to ROS-producing xenobiotics. It also emphasizes the indispensability of rapid *de novo* GSH synthesis during such incidents, something the thioredoxin system is unable to do, which suggests that the thioredoxin system is the “low capacity” leg of antioxidant defense, largely responsible for removing H_2_O_2_ under normal physiological conditions of low oxidative stress, followed by the medium-capacity GSH-reliant system, followed finally by catalase. Simulations indicate that a very important function of catalase is to keep H_2_O_2_ concentration from rising to levels causing necrosis (*>* 500 *µM* [52]) during incidents of extreme oxidative stress; apoptosis is in general more desirable, as it does not provoke an inflammatory response that can damage surrounding tissues.

The most important results of this work from a toxicology perspective are as follows. (1) It was necessary to create a model of endogenous H_2_O_2_ production and removal in hepatocytes as a preliminary to developing a PBPK/PD model that is able to do what was demonstrated here. Neither the peak concentration nor the area under the curve obtained from a PK time course of intracellular DMA*^III^* can be used (at least, not except very crudely) to predict whether a given dose/exposure concentration of DMA*^III^* (or a given oral dose of DMA*^V^*) will result in hepatocyte cell death by the time the arsenical is eliminated, and, if so, when cell death will commence and to what extent it will occur. One must solve the model equations presented here and determine whether or not a critical time point, call it *t_crit_*, will ever arrive, beyond which GSH will be too depleted for GPX (requiring GSH as cofactor to remove H_2_O_2_) to effectively counteract the elevated rate of H_2_O_2_ production associated with the remaining DMA*^III^* PK time course. (2) Existence of an exposure concentration threshold for DMA*^III^* in medium to cause cell death during *in vitro* exposure: Although the function *f* ([DMA*^III^*]) identified from Naranmandura and coworkers’ data indicates no intracellular DMA*^III^* concentration below which this xenobiotic does not elevate the rate at which H_2_O_2_ is produced (i.e., it does not have a sigmoidal shape), the need for GSH to be critically depleted before cell death can occur explains the existence, in Naranmandura’s experimental data, of a threshold concentration for DMA*^III^* in medium to cause detectable cell death within the 24-hr exposure period examined (this threshold is between 0.1 and 1 *µM* in the non-NAC-treated cells, and between 7.5 and 10 *µM* in the NAC-treated cells). (3) In designing *in vitro* experiments in which cells are exposed to various concentrations of a xenobiotic in medium, it is important to choose exposure concentrations that translate into intracellular concentrations similar to those predicted (using *in vivo* PBPK models) for dosing of the living animal in the range of interest. In this work, *f* ([DMA*^III^*]) was identified from data reported for experiments in which the tested DMA*^III^* exposure concentrations are only expected to translate (according to the predictions of the *in vitro* PBPK model presented here) into intracellular DMA*^III^* concentrations *<* 30 *µM*. In contrast, the 2019 Bilinsky PBPK model predicts that oral dosing of mouse with its LD50 for DMA*^V^* results in DMA*^III^* concentrations in liver ten times this value. *f* almost certainly cannot be accurately extrapolated out this far, especially given its super-quadratic term, which may level off at some sufficiently high DMA*^III^* concentration. That is why the model predicts a H_2_O_2_ concentration in mouse liver *∼* 1 *M* for this dose, too high to be physiologically realistic. To realistically model the consequences for liver of the oral LD50 dose of DMA*^V^*, experiments analogous to Naranmandura’s and coworkers’ should be performed using higher DMA*^III^* exposure concentrations, translating into intracellular DMA*^III^* concentrations *∼* 100 *µM*, with cell mortality being assessed after a much shorter time interval than 24 hours. This would enable *f* ([DMA*^III^*]) to be accurately identified over the relevant domain (that is, range of intracellular DMA*^III^* concentration). (4) In their 2020 meta-analysis of studies investigating the effects of arsenic exposure on cellular GSH metabolism, Ran and coworkers reported that *in vitro* preparations of cells chronically exposed to low-dose arsenic show activation of the p38/Nrf2 pathway (indicative of oxidative stress) and, incident to this, upregulated GCLC (the catalytic subunit of GCL) expression and an increased capacity to synthesize GSH. [47]. The insights yielded by this work indicate that these metabolic alterations would make it more difficult for a cell to achieve an apoptotic milieu, both by working against cellular attempts to deplete GSH via its active extrusion, and by keeping H_2_O_2_ concentration below the range needed for a role in the apoptotic mechanism. This is consistent with arsenicals’ known role as carcinogens in chronic, low-dose exposure scenarios. That these metabolic alterations are caused by oxidative stress-triggered pathways such as Nrf2 suggests, on theoretical grounds related to point (2) above, that a nonlinear rather than “linear no-threshold” dose-response relationship exists between arsenical exposure and increased cancer risk, consistent with Lamm and coworkers’ 2021 epidemiological review [34].

## Appendix A. Computational models and rationale

### Appendix A.1. in vitro computational model of endogenous H_2_O_2_ production and removal in rat hepatocytes in cell culture (no xenobiotic present)

I use separate compartments for mitochondria and cytoplasm to allow for greater detail in modeling the production and neutralization of H_2_O_2_, which is generated by mitochondria as well as by organelles of the cytoplasm. I also do this because GSH synthesis is confined to the cytoplasm, hence only cytoplasmic GSH concentration is relevant to product inhibition on GCL. I model the mitochondrial compartment as occupying 15% of the cell volume [24], and the cytoplasmic compartment (including cytoplasmic organelles) as occupying the rest. I now present and explain the equations of my model of endogenous H_2_O_2_ production and removal in rat hepatocytes. Nonlinear kinetic terms (indicated by function notation) are given at the end of this section, except where they are of the well-known Michaelis-Menten form.

The equations governing the concentrations of H_2_O_2_ in the mitochondrial and cytoplasmic compartments are given below.

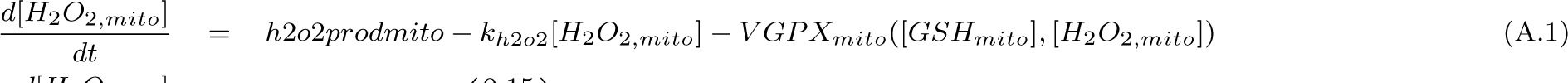

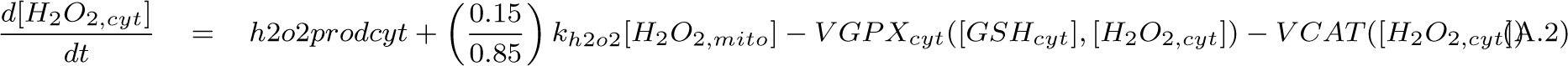

In mitochondria, stray electrons falling off the electron transport chain combine with oxygen to form superoxide, which is then converted to hydrogen peroxide (H_2_O_2_) by superoxide dismutase. I do not model the details of this process but assume a constant rate of production of H_2_O_2_ in mitochondria. Some of the hydrogen peroxide produced diffuses into the cytoplasm, and the rest is neutralized by mitochondrial glutathione peroxidase (GPX), which combines two molecules GSH with one molecule H_2_O_2_ to form glutathione disulfide (GSSG). H_2_O_2_ diffusing from the mitochondria appears in the cytoplasm, where H_2_O_2_ is also locally produced by organelles including the endoplasmic reticulum [8]. Cytoplasmic H_2_O_2_ is neutralized to GSSG by cytoplasmic GPX. I emphasize that in this work, what is called “glutathione peroxidase” and symbolized by GPX actually represents the action of all antioxidant enzymes serving to reduce either H_2_O_2_ itself, or proteins oxidized by H_2_O_2_, and ultimately reliant upon the cofactor GSH for their reduction back to active form (see Introduction).

When determining values for the parameters *h*2*o*2*prodcyt*, *h*2*o*2*prodmito*, and *k_h_*_2*o*2_, I assumed that H_2_O_2_ has normal physiological concentration 0.005 *µM* in mitochondria and 0.01 *µM* in cytoplasm [8]. H_2_O_2_ is produced in liver at rate 50 nmol/min/g tissue [58] and Boveris et al. state that, in rat liver, H_2_O_2_ diffusing from the mitochondria accounts for 40% of H_2_O_2_ appearing in cytoplasm, with the other 60% produced by organelles including endoplasmic reticulum [8]. This enables us to determine *h*2*o*2*prodcyt* and *k_h_*_2*o*2_. Table 2 of [9] enabled us to estimate an efflux of H_2_O_2_ from isolated mitochondria of 0.4 nmol/min/mg protein under normal physiological conditions. Treburg et al. [66] state that such measurements underestimate the rate of production of H_2_O_2_ inside the mitochondrial matrix, due to its consumption there by mitochondrial glutathione peroxidases, and they provide a formula relating the true rate of production to the mitochondrial efflux. Using this formula I obtain 1 nmol/min/mg protein for the true rate of production, 2.5 times higher than the measured efflux. Hence, I assume that *h*2*o*2*prodmito* is 2.5 times the rate needed to balance diffusion into cytoplasm. *V_max_* values for the cytoplasmic and mitochondrial GPX reactions were then chosen to achieve balance (right-hand sides of the differential equations equal 0 when concentration of H_2_O_2_ is 0.005 *µM* in mitochondria and 0.01 *µM* in cytoplasm).

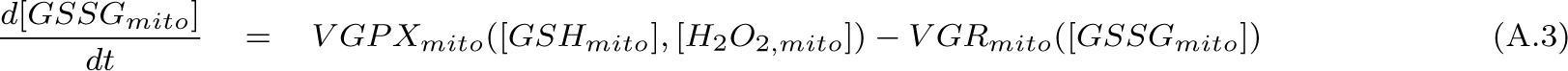

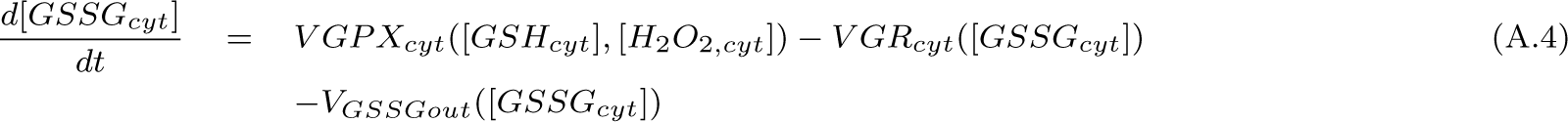

The equations governing the concentrations of GSSG in the mitochondrial and cytoplasmic compartments are given below.

GSSG cannot directly exit the mitochondria [36] but must first be reduced by glutathione reductase (GR) to GSH. Hence, *V_max_*for the mitochondrial GR reaction was chosen to balance the velocity of the mitochondrial GPX reaction when substrates have their steady-state values (see Table 1). GSSG in cytoplasm is exported through the apical membrane (*in vivo* to bile) at rate 0.4 *nmol/min/gram* liver [4]. I convert this to the units (*µM/hr*) by first converting to *µmol/hr/kg* liver, then multiplying by the factor 1*kg/*0.7*L* under the assumption that the liver is 70% water by weight, and finally dividing by 0.85, the fraction of liver volume occupied by the cytoplasm. This yields 40 *µM/hr* for the rate of loss of cytoplasmic GSSG due to export. Export has Michaelis-Menten kinetics with *K_m_*=400 *µM* [3]. *V_max_* was chosen to yield agreement with the 40 *µM/hr* figure when cytoplasmic GSSG has its steady-state concentration. *V_max_* for the cytoplasmic GR reaction was then chosen to achieve balance.

The equations governing the concentrations of GSH in the mitochondrial and cytoplasmic compartments are given below.

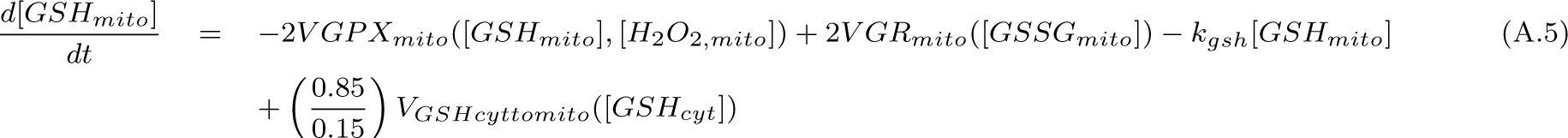

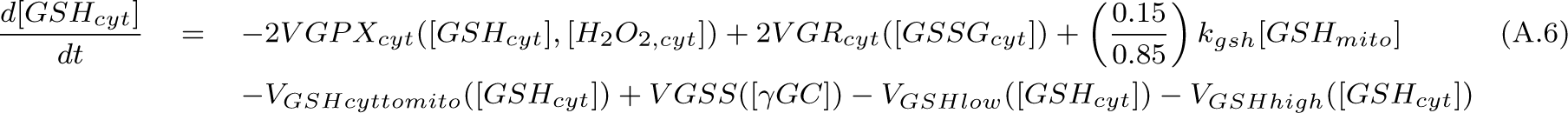

GSH is synthesized only in the cytoplasm, in a sequence of two reactions. In the first reaction, catalyzed by glutamate-cysteine ligase (GCL), the amino acids cysteine and glutamate are combined to form *γ*-glutamylcysteine. Then, *γ*-glutamylcysteine is combined with glycine to form GSH, in a reaction catalyzed by glutathione synthetase (GSS). In this work I do not model glutamate or glycine concentration dynamics, instead holding them constant at their normal physiological values; for the purposes of this model this is an acceptable approximation. GSH is exported through the sinusoidal membrane (*in vivo* to blood plasma) via low- and high-affinity transporters [50]. I use the same kinetic forms and *K_m_* values for these transporters as in Reed et al. [50], and choose *V_max_* values such that, when GSH production is blocked, cytoplasmic GSH declines with the experimentally determined half-life of 2-3 hours. Following [50], the high-affinity transporter is assumed to have Michaelis-Menten kinetics, and the low-affinity transporter sigmoidal kinetics, given by Eq. A.11. *V_max_* for the GSS reaction was then chosen to achieve balance.

Although only synthesized in the cytoplasm, GSH is exchanged between the cytoplasm and the mitochondria in a dynamic equilibrium. GSH is transported into mitochondria via a high-affinity and a low-affinity transporter, each obeying Michaelis-Menten kinetics [15]. I used data provided in [15] to estimate the *K_m_* values and the ratio of *V_max_*values for the two transporters. I then scaled the *V_max_* values (reported in units of *nmol/sec/mg* microsomal protein) so that GSH is transported from the cytoplasm into mitochondria at the rate measured by Fernandez-Checa et al. [20], 18 *nmol/min/gram* liver (1815 *µM/hr* in the model’s units; see earlier paragraph on cytoplasmic GSSG for a description of how this conversion is performed). Transport of GSH from mitochondria to cytoplasm was assumed to obey linear (first-order) kinetics, with rate constant chosen to balance.

The equation for *γ−*glutamylcysteine is given below.

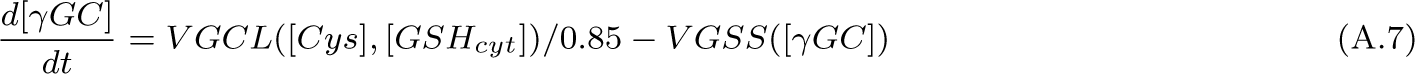

*γ−*glutamylcysteine is produced by glutamate-cysteine ligase (GCL) and consumed by glutathione synthetase (GSS) in the production of GSH. For simplicity in writing the equation for intracellular cysteine (see Eq. A.8), *V GCL* represents the rate at which the whole-cell concentration of cysteine falls due to the consumption of cysteine by the GCL reaction. Since *γ−*glutamylcysteine is produced only in the cytoplasm, *V GCL* must be scaled by 0.85 (the fraction of cellular volume occupied by the cytoplasm) to yield the increase there in the concentration of *γ−*glutamylcysteine. *V_max_* for the GCL reaction was then chosen to achieve balance.

The equation for intracellular cysteine is given by Eq. A.8. This is the first equation where one must specify the environment in which the hepatocytes are situated, in order to model their uptake of cysteine. I use this model as a starting point in modeling the experiments conducted by Naranmandura et al. [43], in which *in vitro* preparations of rat hepatocytes were exposed to various concentrations of DMA*^III^* in medium. Hence, I choose to model the rat hepatocytes as existing in culture and taking up cysteine from medium; in the *in vivo* computational model for mouse presented later in this work, the hepatocytes take up cysteine from the liver capillary bed. In the experimental set-up considered by Naranmandura et al. 20,000 rat hepatocytes are situated in a 100 *µ*L well. I approximate the volume of medium, *vmed*, as the volume of the well. The volume of cell culture, *vcell*, was computed by multiplying 20,000 by the volume of a rat hepatocyte (*≈* 7000 *µm*^3^ [40]) and converting to liters (see Table A.4).

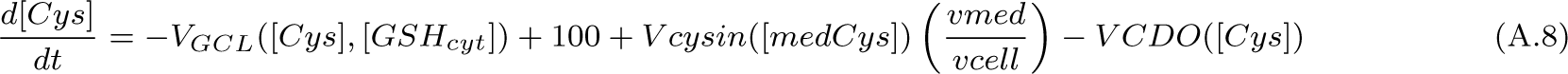

Intracellular cysteine is either consumed by the GCL reaction or catabolized via cysteine dioxygenase (CDO) to sulfate and taurine. At lower concentrations of intracellular cysteine, 90% of cysteine consumption is through the GCL reaction and 10% through the CDO reaction; at high concentrations, 20% of cysteine consumption is through the GCL reaction and 80% through the CDO reaction [61]. I use a quadratic expression to model the kinetics of the CDO reaction (see Eq. A.12), with parameters *a* and *b* chosen so that the relative fluxes through the GCL and CDO reactions at low and high cysteine concentration agree with the above facts. These reactions are mainly fed by the uptake of extracellular cysteine via the ASC transporter. The term *V cysin*([*medCys*]) describes the rate of decrease in medium cysteine concentration (in *µM/hr*) due to this uptake; kinetics are Michaelis-Menten with *K_m_*=2100 *µM* [33]. This term must be multiplied by *vmed* and divided by *vcell* to yield the rate of increase in intracellular cysteine concentration due to uptake. Cysteine is also generated intracellularly, from methionine. I do not model this in detail but assume methionine enters the transulfuration pathway to form cysteine at rate 100 *µM/hr* [50]. *V_max_* for *V cysin* was chosen so that cysteine taken up from medium appears in hepatocytes at rate 1840 *µM/hr*, the rate needed to achieve balance.

The equations governing medium cysteine (medCys) and medium cystine (medCyss) are given below. Cystine is the oxidized form of cysteine. Although hepatocytes cannot usually take up cystine, a dynamic equilibrium exists in medium between the two forms, hence I model both.

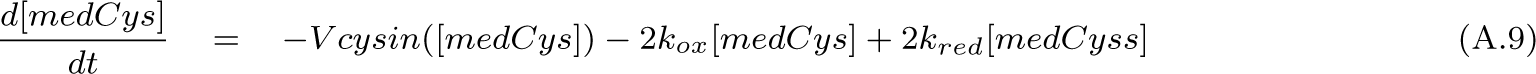

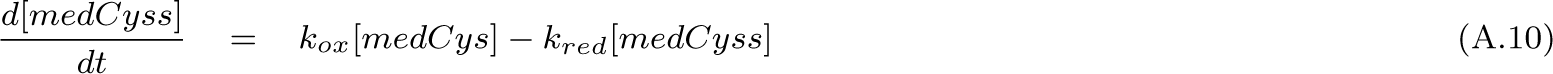

In Naranmandura’s experiments, hepatocytes were cultured with Dulbecco’s Modified Eagle Medium. Dulbecco’s Modified Eagle Medium is formulated with approximately 260.5 *µM* added cystine and no cysteine, but an equilibrium is rapidly reached in which some cysteine is present. Values for *k_ox_* and *k_red_* were chosen so that, when this equilibrium is reached, approximately 20 *µM* cysteine is present, a value which falls within the range of concentrations reported in the literature for cysteine in rat blood plasma under normal physiological conditions (see Table 1).

A note on how I simulate the situation of hepatocytes *in vitro*: cysteine that is taken up from medium by cells is either catabolized via cysteine dioxygenase or used to synthesize GSH. Although GSH is exported, the medium lacks the enzymes present *in vivo* that serve to break down GSH and re-release cysteine into blood. Hence, there is a slow decline of cysteine concentration in medium. In order to compute a steady-state solution (which requires substrate concentrations in medium to be constant) for comparison with normal physiological concentrations in rat hepatocytes, I held the concentration of cysteine in medium constant at 20 *µM* (the normal concentration in rat blood plasma) and solved the model equations. The result is what is given in Table 1.

Equations defining nonlinear kinetic terms appearing in the model are given below. For each term, Table A.3 gives a description and units (column one), the symbol (column two), a reference to the equation defining the term (column three), a list of parameters appearing in the kinetic term (column four), and parameter values with units (column five). Values for parameters not appearing in a nonlinear kinetic term (such as *vcell* and *k_h_*_2*o*2_) are listed without units in column five; units for these parameters are specified in column one. When a nonlinear kinetic term features *K_m_* values for more than one substrate, I write *K_x_* for each of them, where *x* is the substrate; see Eq. A.16 for an example.

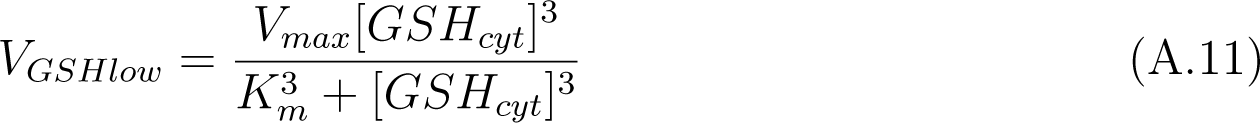

See the discussion of Eq. A.6.

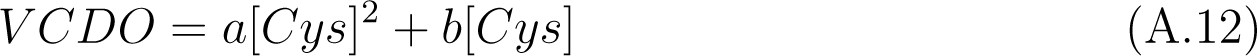

See the discussion of Eq. A.8.

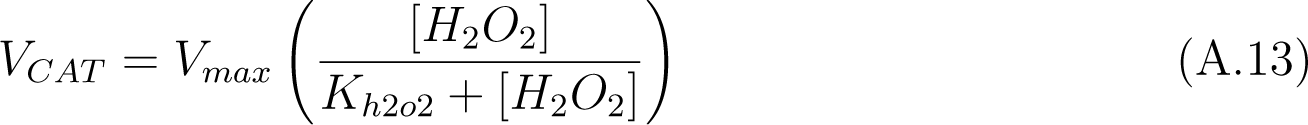

The kinetic form used for catalase is Michaelis-Menten in *H*_2_*O*_2_. I use a *K_m_* for *H*_2_*O*_2_ of 32,300 *µM* [13].

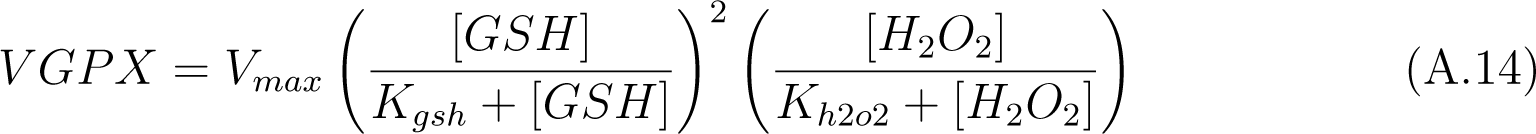

The kinetic form used for glutathione peroxidase is Michaelis-Menten in *H*_2_*O*_2_ and in each of the two required molecules of GSH. I use a *K_m_* for GSH of 1330 *µM* [10] and a *K_m_* for H_2_O_2_ of 6.8 *µM* [71]. In this model, the mitochondrial and cytoplasmic GPX reactions have different *V_max_* values.

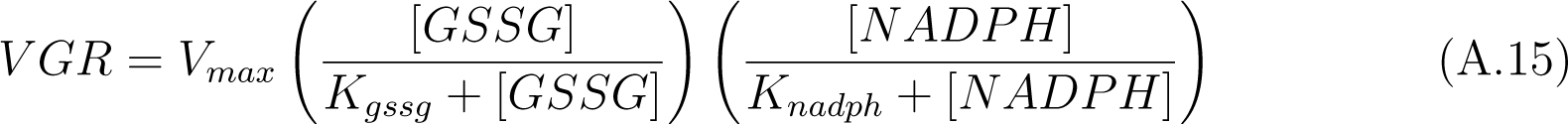

The kinetic form and *K_m_* values for glutathione reductase are taken from [50]. It is Michaelis-Menten in GSSG and in the electron-donor nicotinamide adenine dinucleotide phosphate (NADPH). I do not model NADPH dynamics, but assume it is constant at concentration 50 *µM* [50]. In this model, the mitochondrial and cytoplasmic GR reactions have different *V_max_* values.

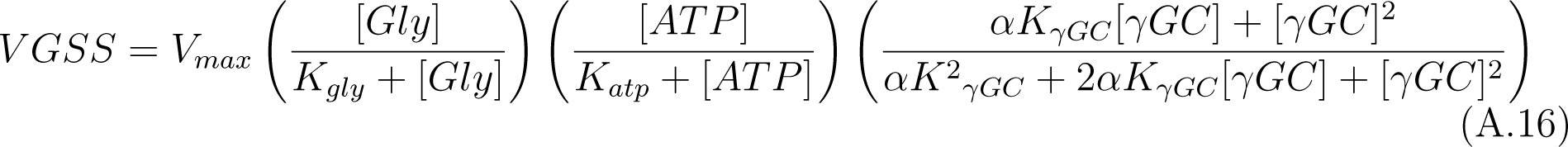

The kinetics of the GSS reaction are taken from [35]. The parameter *α* is a constant known as the “interaction factor”. The value I use for *K_γGC_* was actually obtained for the compound *γ*-glutamyl-*α*-aminobutyrate (gluABA), a common substitute for *γ*-glutamylcysteine in kinetic studies, which does not undergo the complication of thiol oxidation. I do not model glycine dynamics, but assume it is present at concentration [Gly]=3000 *µM* [64]. This reaction also depends on ATP. I do not model its dynamics, but assume it is present at concentration 3000 *µM* [57].

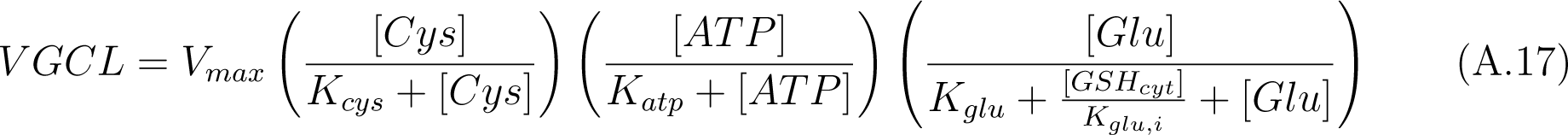

The kinetic form and parameter values for the GCL reaction are taken from [27, 23, 39]. The reaction kinetics feature competitive inhibition of glutamate by GSH. I do not model glutamate dynamics, but assume it is present at constant concentration [Glu]=2000 *µM* [64].

### Appendix A.2. in vitro PBPK/PD model predicting H_2_O_2_ concentration in rat hepatocytes and cell death after exposure to the ROS-producing arsenical DMA^III^

To simulate the experiments carried out by Naranmandura et al., in which rat hepatocytes in culture were exposed to various concentrations of DMA*^III^* in medium for 24 hours and cell death assayed [43], I first must augment my model of endogenous H_2_O_2_ production and removal with an *in vitro*-level pharmacokinetic model for DMA*^III^* and DMA*^V^*. Intracellular DMA*^III^* undergoes oxidation to DMA*^V^* and is released primarily in pentavalent form. In my and collaborators’ previously published PBPK model for oral or intravenous dosing of mice with DMA*^III^* or DMA*^V^* [7], we used diffusion-limited kinetics and obtained permeability and partition coefficients for both DMA*^III^* and DMA*^V^* in liver by fitting to PK data sets. We also obtained a rate constant for the oxidation in liver of DMA*^III^* to DMA*^V^*. Here I scale the rate constant for oxidation and the permeability coefficients so as to model pharmacokinetics in an *in vitro* preparation consisting of 20,000 cells per well, the experimental setup used by Naranmandura et al. Partition coefficients do not need to be scaled. The four PBPK equations added to the rat hepatocyte model are given below. I note that mouse and rat hepatocytes cannot be expected to respond identically to DMA*^III^*. However, in the absence of analogous experiments for mouse hepatocytes, this work serves as a proof of concept.

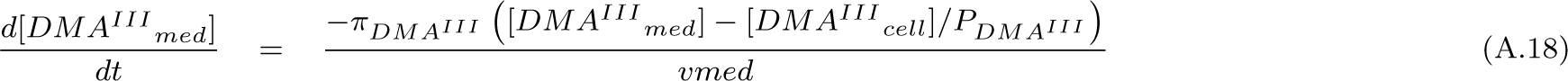

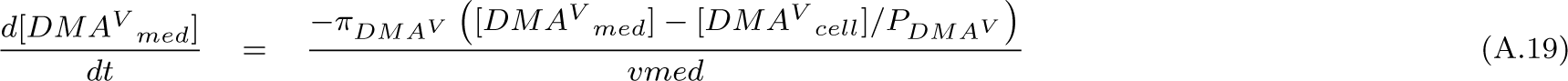

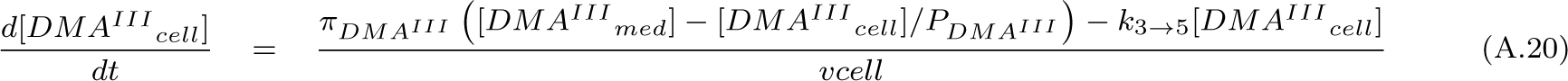

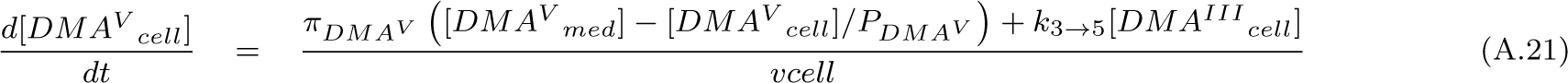

I now model the action of DMA*^III^* on hepatocytes. I alter Eq. A.2 for H_2_O_2_ concentration in cytoplasm, adding a term *f* ([DMA*^III^*]) which models the excess production of H_2_O_2_ caused by DMA*^III^*.

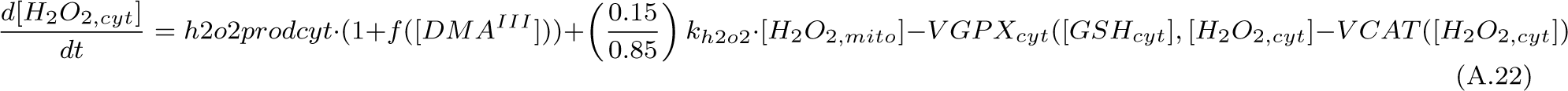

I assume that the per-cell death rate of hepatocytes increases linearly with average intra-cellular H_2_O_2_ concentration until a maximal concentration *h*2*o*2*lim* is reached; beyond this, the cellular death rate is maximal and constant; for a discussion of this H_2_O_2_ ceiling, see section 3.5.1. Below the ceiling, the equation governing the time-evolution of the number *N* of surviving hepatocytes in culture is:

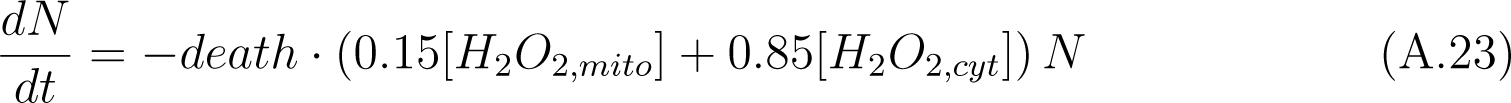

In the above, *death* is the parameter relating H_2_O_2_ concentration to the per-cell death rate, and 0.15[*H*_2_*O*_2,*mito*_] + 0.85[*H*_2_*O*_2,*cyt*_] is the intracellular H_2_O_2_ concentration averaged over the cytoplasmic and mitochondrial compartments. I neglect cellular replication under these distressed circumstances. When this average H_2_O_2_ concentration exceeds *h*2*o*2*lim*, Eq. A.23 is replaced with

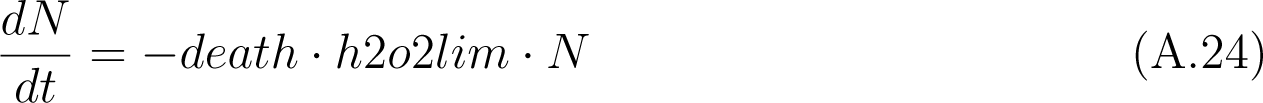

Since I am modeling a situation in which significant cell death occurs, and only living hepatocytes take up cysteine, I must modify Eq. A.9, multiplying the term *V cysin*([*medCys*]) by *fracsurv*, the fraction of surviving hepatocytes. The modified equation is given below.

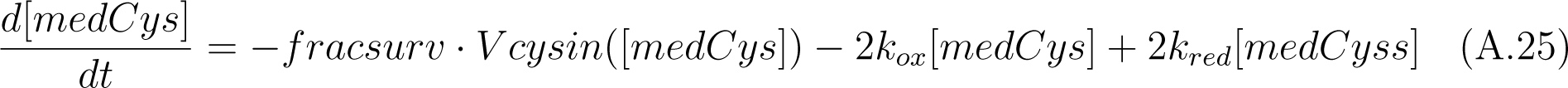

Eq. A.8 remains unaltered because *fracsurv* would multiply both the numerator and denominator of the cysteine uptake term, resulting in cancellation.

### Appendix A.3. Simulation of Naranmandura experiments

Naranmandura et al. cultured hepatocytes in Dulbecco’s Modified Eagle Medium, with or without the addition of 2000 *µM* N-acetylcysteine (NAC), for 24 hours prior to exposing the cells to various concentrations of DMA*^III^*; DMA*^III^* exposure then continued for another 24 hours [43]. Cell viability (fraction of surviving cells) was then assessed. To simulate these experiments, for each of the tested concentrations of DMA*^III^* the following was done. For the case of the non-NAC-treated hepatocytes, initial concentrations of species in hepatocytes and medium were set equal to the values given in Table 1 and the state after 24 hours computed. For the case of the NAC-treated hepatocytes, the same was done, except that the initial concentration of cysteine in medium was set equal to 2020 *µM* (due to the addition of 2000 *µM* NAC). NAC is taken up by hepatocytes and converted to cysteine via acylase; for simplicity, I do not model this process, and treat NAC as cysteine. For these simulations of preliminary culturing, no DMA*^III^* or DMA*^V^* was present. Then, for both the NAC-treated and the non-NAC-treated cells, the concentration of DMA*^III^* in medium was set equal to the tested value (0.1 *µM*, etc.), the solution advanced another 24 hours, and the fraction of surviving hepatocytes computed.

The expression *f* ([DMA*^III^*]), modeling the excess production of H_2_O_2_ in cytoplasm due to the action of DMA*^III^*, and the parameter *death* characterizing the increase in celluar death rate with H_2_O_2_ concentration were determined by fitting to Naranmandura’s data. The functional form arrived at for *f* ([DMA*^III^*]) is given below.

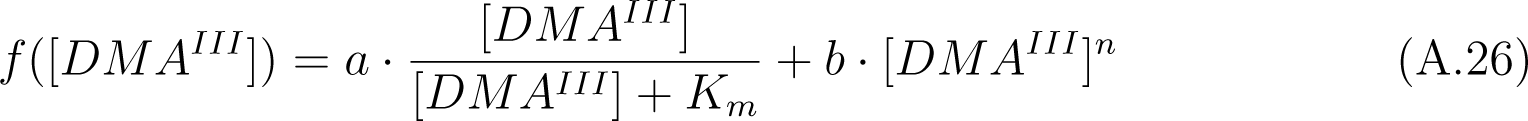

Table A.4 gives all parameter values related to the modeling of DMA*^III^*-induced cytotoxicity.

### Appendix A.4. in vivo computational model of endogenous H_2_O_2_ production and removal in mouse liver (no xenobiotic present)

Before creating a PBPK/PD model of liver injury in mice dosed with DMA*^III^* or DMA*^V^*, I must first create an *in vivo* computational model of endogenous H_2_O_2_ production and removal in the liver of mouse. GSH exported by liver is broken down to its constitutent amino acids by *γ*-glutamyltransferase and dipeptidase located on the membrane of renal proximal tubule [1]. The cysteine which is released can be taken up again by liver. Hence, a cycling of cysteine between liver and blood plasma occurs; for a more extensive discussion of this see Reed et al. [50] and Bilinsky et al. [6]. In the *in vitro* rat hepatocyte model, a decomposition of GSH was not modeled because these enzymes are not present. This *in vivo* computational model also features a dietary input of cysteine to blood plasma.

As in our PBPK model [7], I model the liver tissue proper as well as the plasma portion of the blood occupying the liver capillary bed. The liver takes up cysteine from the capillary bed, and GSH exported by liver initially appears here. I also model (systemic) blood plasma. The hepatic artery and portal vein carry chemical species from the blood plasma to the liver capillary bed; the portal venous flow also carries DMA*^III^* or DMA*^V^* absorbed from an oral dose. The hepatic vein carries species from the capillary bed to the blood plasma. My model features four compartments: blood plasma, the plasma portion of the liver capillary bed, and mitochondrial and cytoplasmic compartments for the liver tissue. For simplicity blood plasma volume in this model is taken to be equal to the total volume of plasma appearing in our previous PBPK model other than that occupying the liver capillary bed. That is: arterial and venous plasma, as well as plasma residing in the capillary beds of the lungs, kidneys, urinary bladder, slowly perfused tissues, and richly perfused tissues.

In addition to intrahepatic concentrations, the *in vivo* computational model computes the concentrations of GSH in blood plasma ([*bGSH*]), GSH in the plasma of the liver capillary bed ([*livcapGSH*]), cysteine in blood plasma ([*bCys*]), cysteine in the plasma of the liver capillary bed ([*livcapCys*]), and cystine in blood plasma ([*bCyss*]). Since cystine is not usually taken up by hepatocytes, I assume it is shunted through the liver capillary bed and do not model cystine in the plasma of the liver capillary bed explicitly.

The five equations associated with the blood plasma and plasma of the liver capillary bed are given below (Eqs. A.27-A.31). *Q_liver_* is the flow in liters per hour of blood plasma through the liver capillary bed. *V_blood_* is the volume of the systemic blood plasma and *V_cap_* is the volume of plasma in the liver capillary bed. *V_tiss_* is the volume of mouse liver tissue. Aside from *V_blood_*, which is calculated as described in the first paragraph of this section, the values of these parameters are as in [7]. The values of parameters appearing in extrahepatic terms, the *V_max_* for uptake of cysteine by liver, and the steady-state values of the concentrations [*livcapCys*] and [*livcapGSH*] fell out of the requirement that the steady-state values of the concentrations [*bGSH*], [*bCys*], and [*bCyss*] be equal to their normal physiological values (see Table 2) and that cysteine appear in liver at the rate needed to feed the GCL reaction. Parameter values are given in Table A.5. I begin with the equation for GSH concentration in blood plasma.

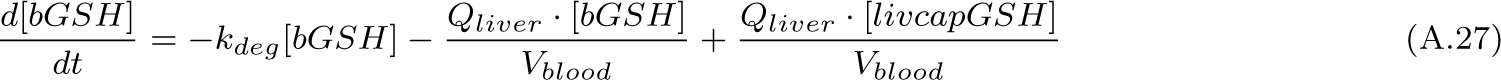

GSH exits the blood plasma and enters the liver capillary bed at rate *Q_liver_ ·* [*bGSH*]. GSH enters the blood plasma from the liver capillary bed at rate *Q_liver_ ·* [*livcapGSH*]. These terms must be divided by *V_blood_* to yield rates of change in concentration. The parameter *k_deg_* dictates the rate at which GSH in blood plasma is broken down by *γ*-glutamyltransferase and dipeptidase to its component amino acids.

The equations for cysteine and cystine concentration in blood plasma are given below.

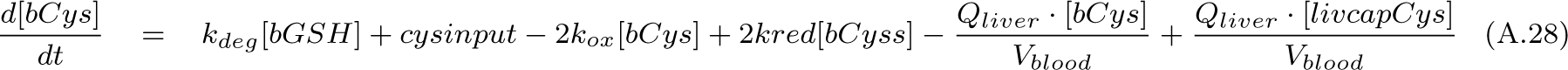

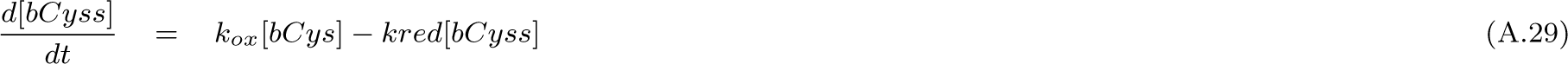

I model the input of cysteine to blood plasma from dietary sources, the appearance of cysteine in blood plasma due to the breakdown of GSH, the exchange of cysteine between the blood plasma and the liver capillary bed, and the interconversion of cysteine and cystine. Dietary cysteine intake in humans is 9 *µmol/hr/kg* body weight [70]; in the absence of mouse-specific data I use this figure. In mouse, this would translate to a rate of appearance in blood plasma of 197 *µM/hr*. This is very close to my value of *cysinput*=170 *µM/hr*, chosen to achieve balance.

The equation for GSH concentration in plasma of the liver capillary bed is given below.

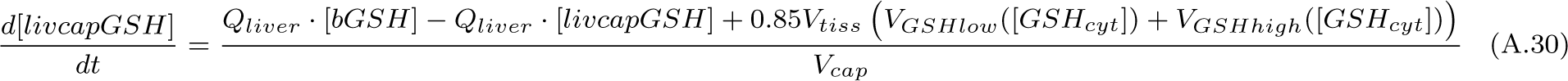

I model the export of GSH from liver via the high- and low-affinity transporters, and the exchange of GSH between blood plasma and the plasma of the liver capillary bed.

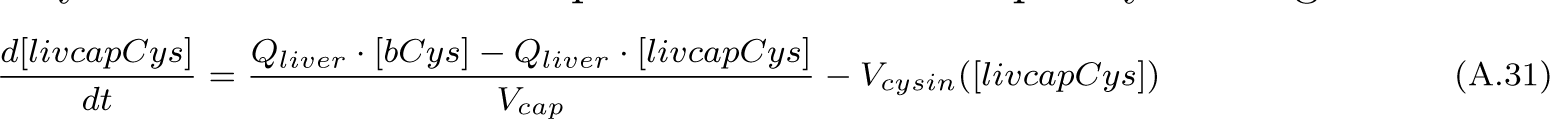

The equation for cysteine concentration in plasma of the liver capillary bed is given below.

I model the uptake of cysteine by liver (units *µM/hr*) and the exchange of cysteine between blood plasma and the plasma of the liver capillary bed.

The intrahepatic equations are identical in form to those for rat hepatocytes, except for the kinetics used for the GCL reaction; see Eq. A.32 below.

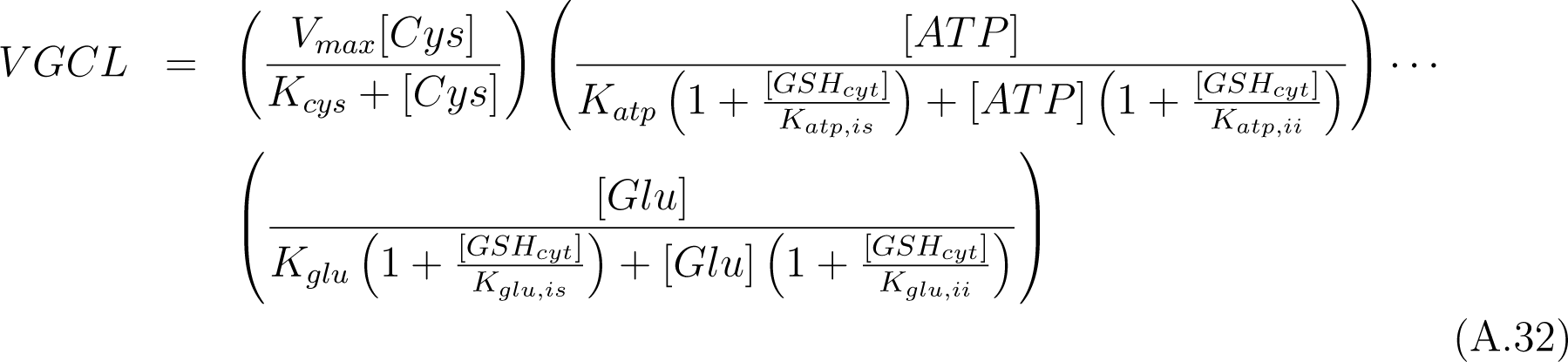

The kinetics for the GCL reaction are taken from Chen et al. [14], who studied the kinetics of GCL in mouse liver. I do not model glutamate or ATP dynamics, but assume they are present at constant concentrations [Glu]=1400 *µM* [37] and [ATP]=3000 *µM* [57]. In the absence of mouse-specific kinetic data, I use the kinetics for the GSS reaction given by Eq. A.16. In computing the velocity of the GSS reaction I assume that glycine is present at constant concentration 1700 *µM*, consistent with measurements in mouse liver by Marquez et al. [37].

Finally, normal physiological concentrations of chemical species in liver are somewhat different in mouse than in rat. Since these concentrations are input arguments for computing fluxes, *V_max_* values in Table A.5 are in general different from those in Table A.3 for the same reactions, as are the parameters *a* and *b* used in computing the velocity of the CDO reaction.

### Appendix A.5. in vivo PBPK/PD model for intravenous or oral dosing of mouse with DMA^III^ or DMA^V^

I conjoin my and coworkers’ 2019 PBPK model for oral or intravenous dosing of the mouse with DMA*^III^* or DMA*^V^* [7] with the *in vivo* computational model presented here for endogenous H_2_O_2_ production and removal in the liver of mouse to form an *in vivo* mouse PBPK/PD model for these arsenicals. One of the predictions of our 2019 PBPK model is that orally ingested DMA*^V^* undergoes extensive reduction in the gastrointestinal tract prior to entering the bloodstream; this has now been experimentally confirmed [67]. Hence, some of an oral dose of DMA*^V^* enters the liver as DMA*^III^*. As was done in Eq. A.22, I alter the term for H_2_O_2_ production by organelles of the cytoplasm to include a term *f* ([DMA*^III^*]) describing increased production of H_2_O_2_ due to the presence of DMA*^III^*. I also include Eq. A.23, modeling hepatocyte death, and scale *V_tiss_*in Eq. A.30 and *V_cysin_*in Eq. A.31 by *fracsurv*, the fraction of surviving hepatocytes.

In this first model, I assume that DMA*^III^* acts upon mouse hepatocytes and rat hepatocytes in the same way, and use the parameter values appearing in Table A.4 in constructing the *in vivo* mouse model for the action of this arsenical. I note that a ceiling concentration of 150 *µM* H_2_O_2_ was chosen, above which cellular death rate is assumed to be maximal and constant. This was necessary because the current model overpredicts intracellular H_2_O_2_ concentration for the scenario of oral dosing of mouse with 650 mg As/kg DMA*^V^* (the oral LD50 [54]), a scenario I wish to simulate here as a proof of concept, due to the application of the function *f* ([DMA*^III^*]), calibrated from Naranmandura’s *in vitro* data (in which the maximum intracellular DMA*^III^* was about 25 *µM*, for the highest exposure level). In the scenario of oral dosing of mouse with its DMA*^V^* LD50, hepatic intracellular DMA*^III^* is pre-dicted to an order of magnitude higher, peaking near 250 *µM*. Not using a ceiling results in all hepatocytes dying rapidly. Hence, a ceiling was used; the choice of 150 *µM* H_2_O_2_ results in a situation in which the LD50 dose causes the death of 2/3 hepatocytes. This target for hepatocyte death was chosen from the consideration that hepatectomies which are more extensive than this result in morbidity and/or mortality [11]; the origin of this criterion for connecting lethal dose to fraction hepatocytes killed is [5]. The ceiling can be dispensed with (or raised to a more accurate, higher level) if data analogous to Naranmandura’s becomes available for mouse hepatocytes exposed to DMA*^III^* concentrations in medium that translate into intracellular concentrations *∼* 100 *µM* (and possibly assayed for death after a period significantly less than 24 hours).

**Table A.3:**
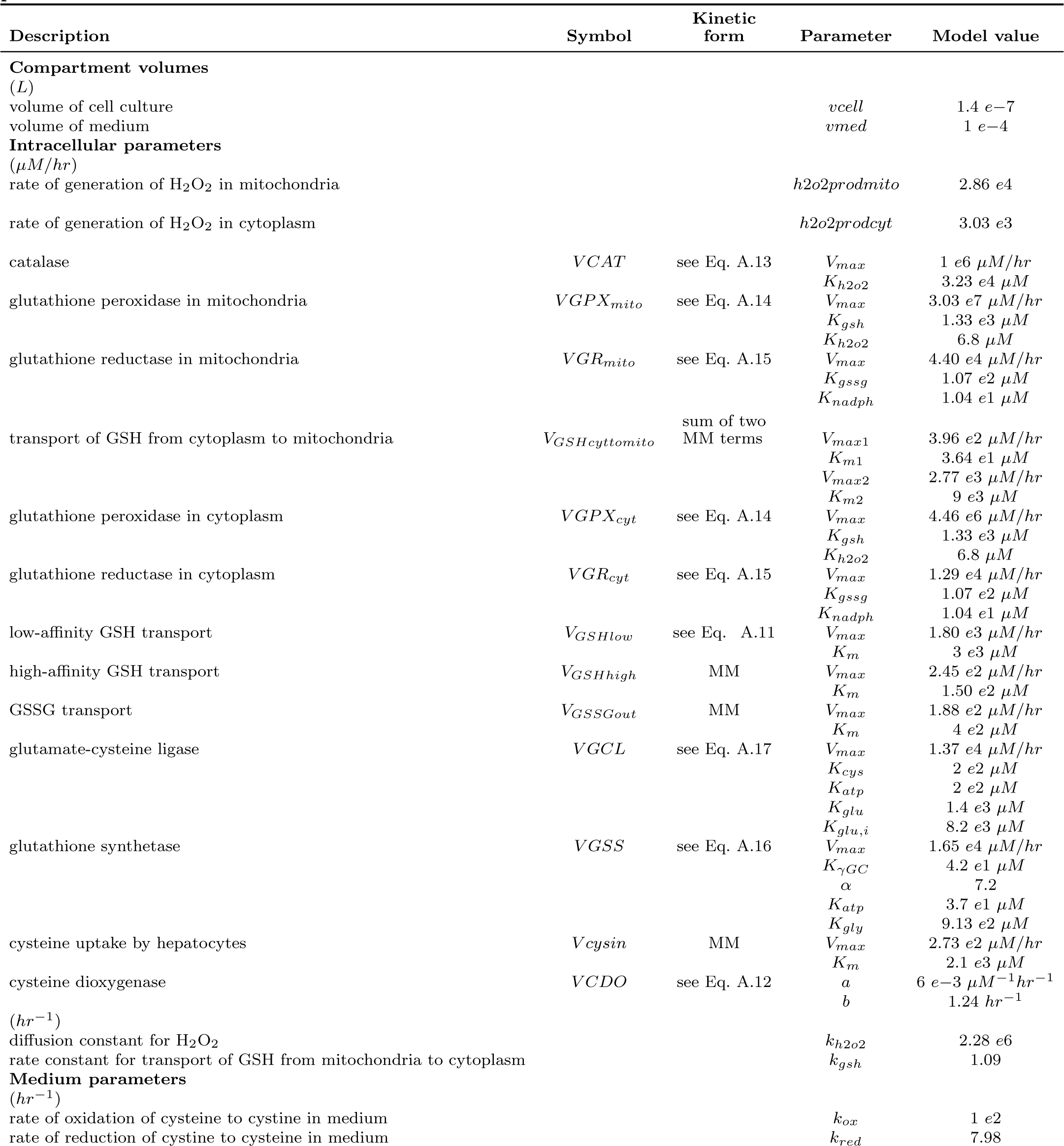
Parameters for the *in vitro* computational model of endogenous H_2_O_2_ production and removal in rat hepatocytes in cell culture subsisting on Dulbecco’s Modified Eagle Medium. Sources and/or rationale for the parameter values appearing in this table are provided in the description of model equations. In general, intrinsic kinetic parameter values (e.g. *K_m_* values) are taken directly from the published literature, while *V_max_* values and other parameters are chosen to result in reaction and transport fluxes at steady-state in agreement with the published literature.

**Table A.4:** Pharmacokinetic and pharmacodynamic parameters for the *in vitro* PBPK/PD model of DMA*^III^* -induced oxidative stress and resultant cell death in rat hepatocytes in cell culture. Partition coefficients and (after scaling to describe the *in vitro* situation) permeability coefficents and the rate of oxidation of DMA*^III^* to DMA*^V^* in hepatocytes are taken from Bilinsky et al. [7]. Pharmacodynamic parameters were obtained as described in section 3.2.1.

| Description | Parameter | Model value |
| --- | --- | --- |
| **Pharmacokinetic parameters** |  |  |
| partition coefficient for DMA<sup>III</sup> | $P_{DMA^{III}}$ | 1 e2 |
| partition coefficient for DMA<sup>V</sup> | $P_{DMA^V}$ | 1 |
| rate of oxidation of DMA<sup>III</sup> to DMA<sup>V</sup> | $k_{3 \rightarrow 5}$ | 9.28 e-7 L/hr |
| permeability coefficient for DMA<sup>III</sup> | $\pi_{DMA^{III}}$ | 2.32 e-6 L/hr |
| permeability coefficient for DMA<sup>V</sup> | $\pi_{DMA^V}$ | 1.86 e-7 L/hr |
| **Parameters for DMA<sup>III</sup>-induced excessive H<sub>2</sub>O<sub>2</sub> production** |  |  |
| | $K_m$ | 0.2 $\mu M$ |
| | $a$ | 2.2 |
| | $b$ | 5 e-4 $\mu M^{-2.5}$ |
| | $n$ | 2.5 |
| **Parameters for H<sub>2</sub>O<sub>2</sub>-associated cell death** |  |  |
| H<sub>2</sub>O<sub>2</sub> concentration at and above which death rate is maximal | $h2o2lim$ | 150 $\mu M$ |
| per-cell death rate of hepatocytes per $\mu M$ H<sub>2</sub>O<sub>2</sub> | $death$ | 9 e-4 $\mu M^{-1} hr^{-1}$ |

**Table A.5:**
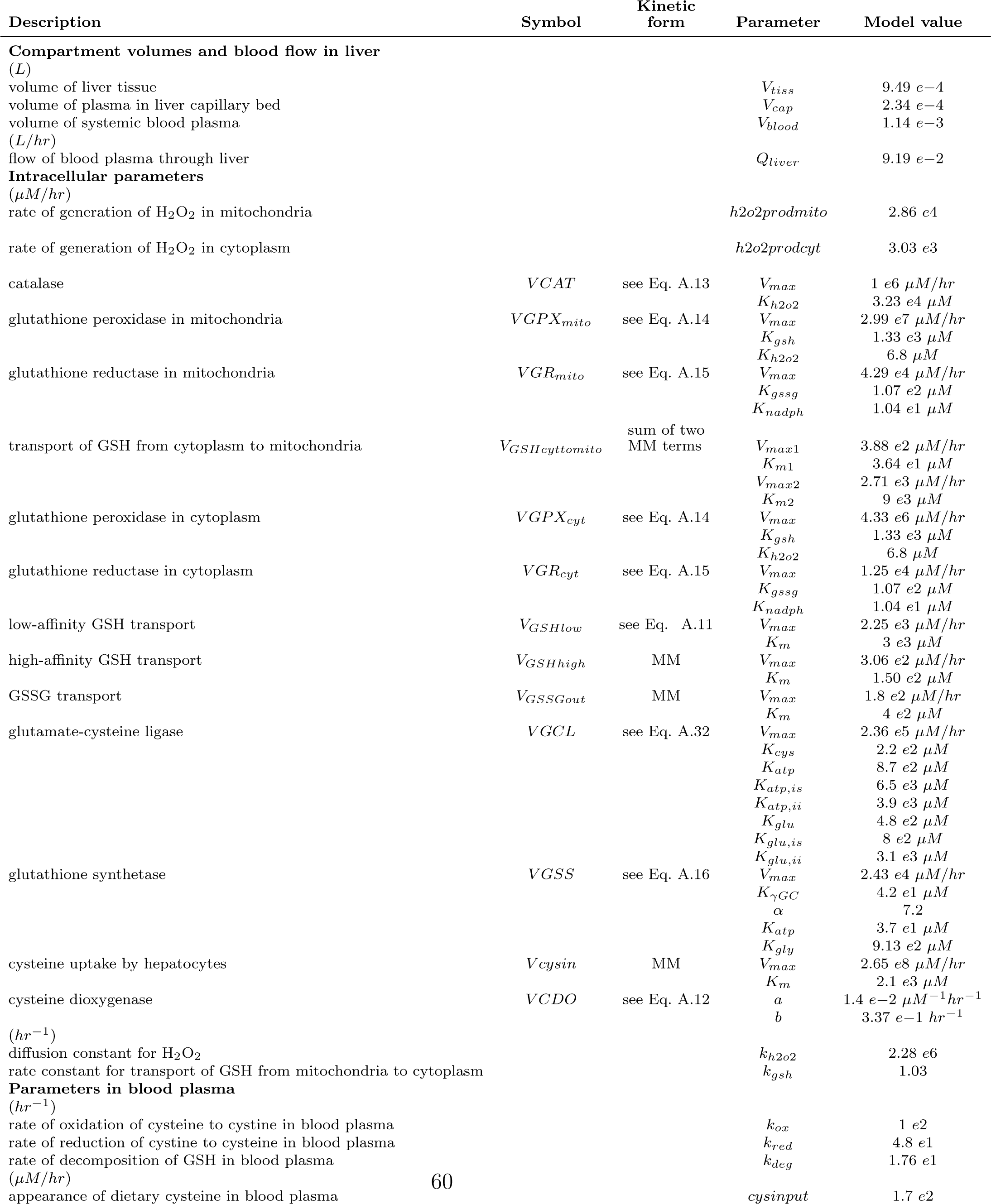
Parameters for the *in vivo* computational model of endogenous H_2_O_2_ production and removal in liver of mouse. Sources and/or rationale for the parameter values appearing in this table are provided in the description of model equations.

## Acknowledgements

This study was funded by an Interagency Agreement between the US Food and Drug Administration (FDA) and the US National Institute of Environmental Health Sciences (NIEHS) (FDA IAG #224-17-0502 and NIEHS IAG #AES12013). This manuscript was reviewed in accordance with FDA procedures prior to submission. This manuscript reflects the views of the authors and does not necessarily reflect those of the FDA or the NIEHS.

## Availability of data and materials

The data set analyzed in this paper and used for model formulation is published in [43]. Model codes will be available on GitHub upon paper publication.

